# Crystal structures of forty- and seventy-one-substitution variants of hydroxynitrile lyase from rubber tree

**DOI:** 10.1101/2025.03.30.646168

**Authors:** Colin T. Pierce, Panhavuth Tan, Lauren R. Greenberg, Meghan E. Walsh, Ke Shi, Alana H. Nguyen, Elyssa L. Meixner, Sharad Sarak, Hideki Aihara, Robert L. Evans, Romas J. Kazlauskas

**Affiliations:** Department of Biochemistry, Molecular Biology and Biophysics, University of Minnesota, Minneapolis, MN 55455, USA

**Keywords:** engineered protein, hydroxynitrile lyase, α/β-hydrolase fold, esterase, SABP2

## Abstract

The α/β-hydrolase fold family contains mostly esterases but includes other enzymes such as hydroxynitrile lyase from *Hevea brasiliensis* (rubber tree, *Hb*HNL). *Hb*HNL shares 44% sequence identity and a Ser-His-Asp catalytic triad with esterase SABP2 (salicylic acid binding protein 2 from *Nicotiana tabacum* (tobacco)). To identify how large a region within *Hb*HNL influences the positions of the catalytic residues, we created variants where increasingly large regions surrounding the substrate-binding site had identical amino acid sequences to those in SABP2. Variant HNL40 contains 40 mutations (two inserted amino acid residues, 38 substitutions), shares 59% sequence identity with SABP2, and is identical in sequence to SABP2 within 10 Å of the substrate-binding site. Variant HNL71 contains 31 additional substitutions for a total of 71 changes (two insertions, 69 substitutions) and shares 71% sequence identity with SABP2. The sequences within 14 Å of the substrate-binding site are identical in SABP2 and HNL71. The crystal structures of HNL40 and HNL71 show that the positions of main chain Cɑ atoms move from their positions in *Hb*HNL to more closely match those in SABP2 (RMSD = 0.51 Å over 235 Cɑ atoms for HNL40, 0.41 Å over 219 Cɑ atoms for HNL71) and even more closely in the region within 10 Å of the substrate-binding site (RMSD = 0.38 Å over 58 Cɑ atoms for HNL40, 0.28 Å over 53 Cɑ atoms for HNL71). The pattern of tunnels in HNL40 and HNL71 are similar to each other and intermediate between the pattern in *Hb*HNL and SABP2.

**Synopsis:** Variants HNL40 and HNL71 of hydroxynitrile lyase from *Hevea brasiliensis* contain 40 and 71 mutations, respectively, to make regions surrounding the substrate-binding site identical in sequence to esterase SABP2. X-ray structures reveal increasing similarities to SABP2 in HNL40 and HNL71.

**PDB reference:** hydroxynitrile lyase from *Hevea brasiliensis* with forty mutations, 8SNI, hydroxynitrile lyase from *Hevea brasiliensis* with seventy-one mutations, 9CLR

## 1. Introduction

*Hb*HNL, hydroxynitrile lyase from *Hevea brasiliensis* (rubber tree), and SABP2, an esterase from tobacco, are homologs with 44% sequence identity (114 identical residues, 146 differing over 260 positions) (Fig. 1). The two proteins have the same α/β-hydrolase protein fold and the same Ser-His-Asp catalytic triad.

**Figure 1.**
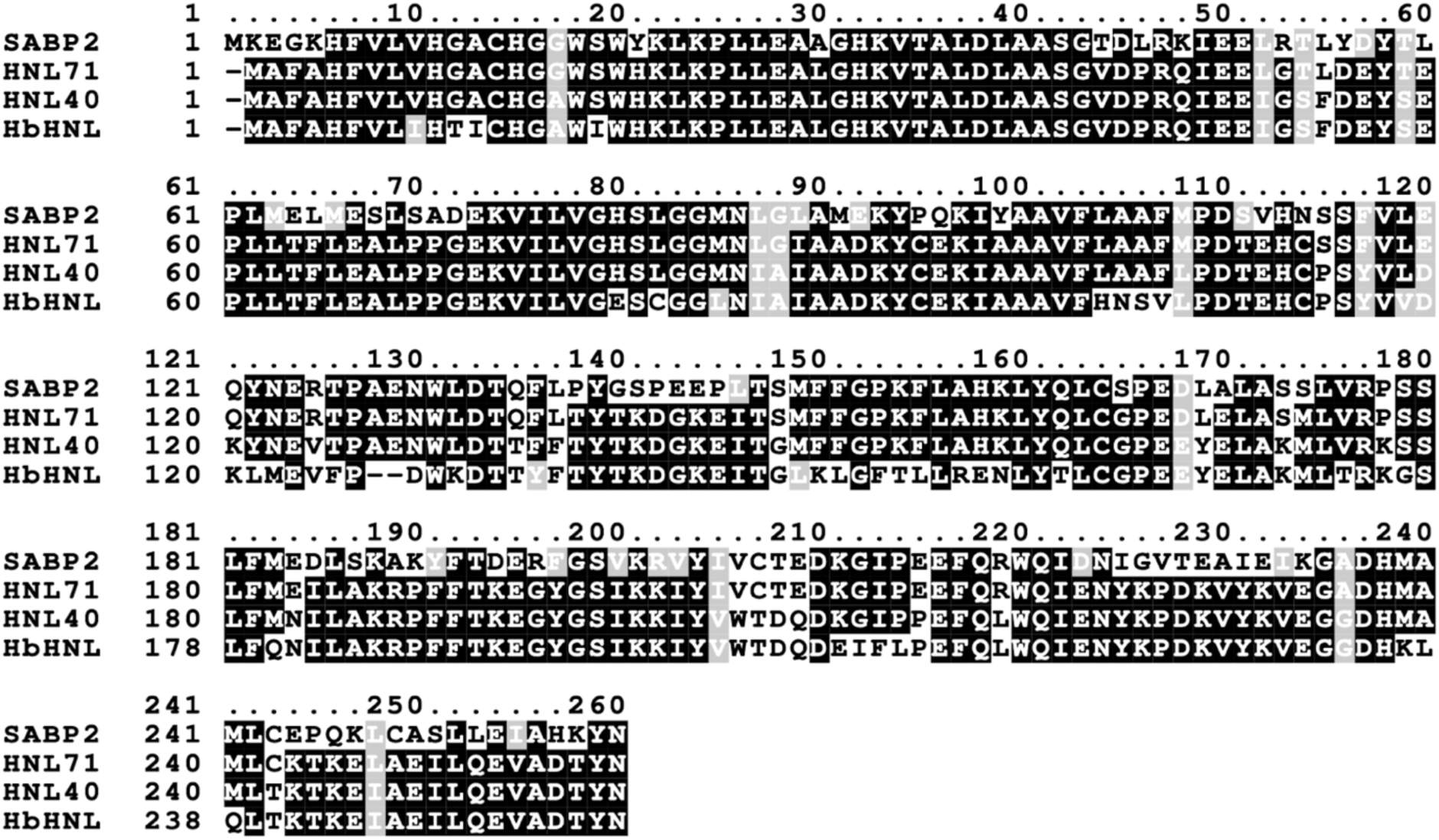
Sequence alignment of HNL40 and HNL71 to the starting sequence *Hb*HNL and the target sequence SABP2. Positions with identical amino acids are shaded black, positions with similar amino acids are shaded gray and positions with dissimilar amino acids are unshaded. The catalytic triad is conserved in all four proteins and occurs at Ser80-His237-Asp209 in the HNL40/71 numbering. Sequences were aligned with Clustal Omega (Sievers et al., 2011) and shaded with BOXSHADE (https://junli.netlify.app/apps/boxshade/). HNL40 and HNL71 also contained an N- or a C-terminal His6 tag, respectively, which are not shown in this alignment

Despite these similarities, both the main chain positions and the positions of the catalytic atoms differ between the two proteins (Fig. 2). The overall Cɑ atom positions in *Hb*HNL differ from those in SABP2 by an average of 0.68 Å for the best 213-218 comparisons out of a total of 256 atoms. Increasing the number of comparison pairs to 249 out of 256 doubles the average difference in Cɑ atom positions to 1.4 Å. The Cɑ atom positions of the catalytic triad (Ser81, His238, Asp210; SABP2 numbering) and oxyanion hole (Ala13, Leu82) differ by an average of 0.8±0.3 Å. Similarly, the five catalytic atoms of the catalytic triad (Ser Oɣ, His Nε, Asp Oδ) and oxyanion hole (OX1 N, OX2 N) differ by a similar amount, an average of 0.8±0.4 Å. We hypothesized that adding substitutions (and insertions) to *Hb*HNL from SABP2 in the regions closest to the substrate-binding region would make the active sites of the *Hb*HNL variants identical to the active site of SABP2.

**Figure 2.**
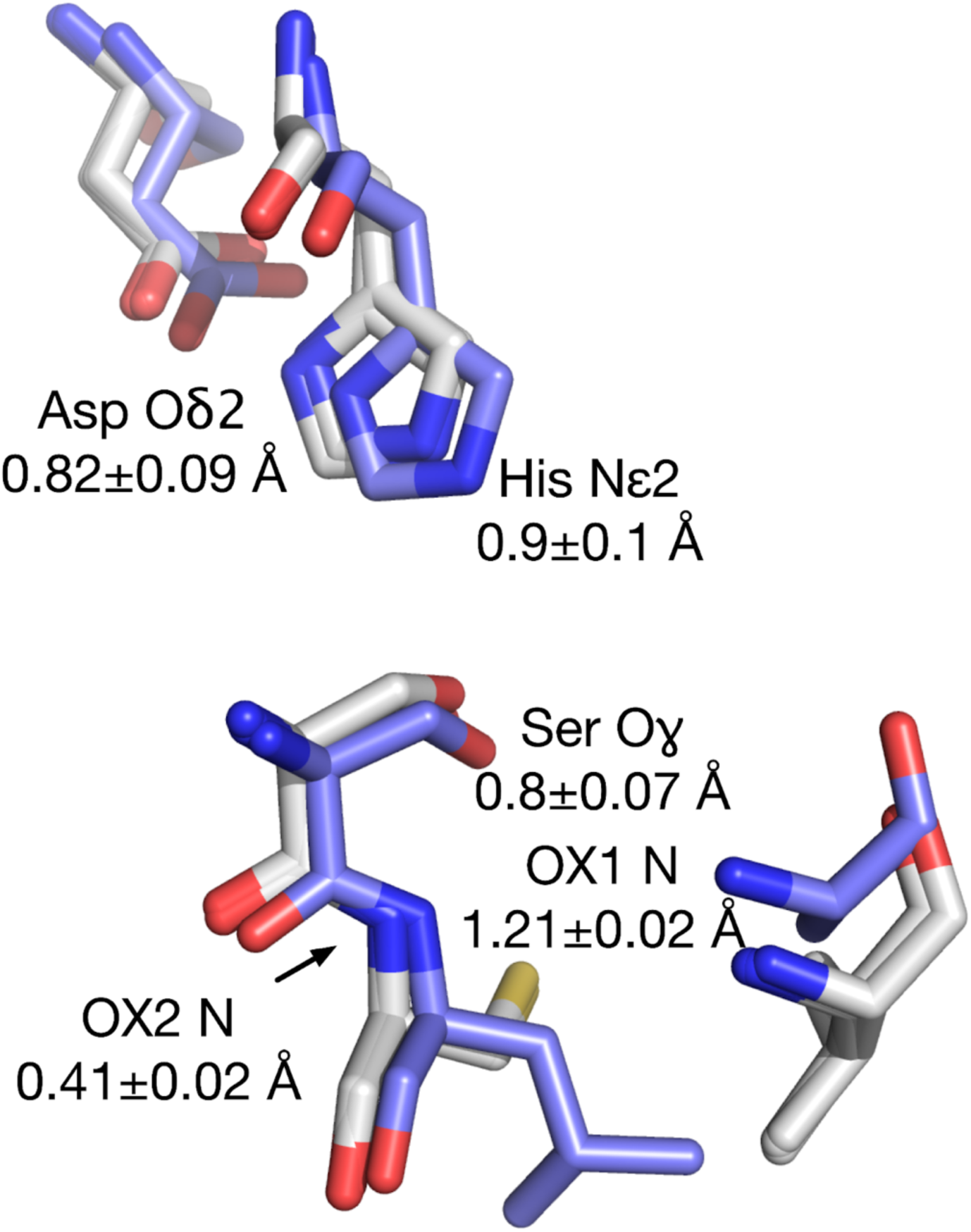
Overlay of the catalytic residues of three ligand-free *Hb*HNL structures (white carbons, pdb entries 6yas, 3c6x, 2g4l) onto the structure of SABP2 (blue carbons, pdb entry 1y7h, contains a small ligand, thiocyanate (not shown) in the active site). The alignment minimized the RMSD of between 213-218 corresponding Cɑ atom pairs in the proteins. The average deviation was 0.68 Å over the aligned Cɑ atoms out of a total of 256. The Cɑ atoms of the catalytic triad (Ser, His, Asp Oδ) and the oxyanion hole (OX1, OX2) deviated by an average Cɑ deviation, an average of 0.8±0.3 Å slightly more than the average Cɑ deviation. The positions of the catalytic atoms identified in the figure deviated by a similar average amount, 0.8±0.4 Å.

HNL40 and HNL71 are variants of HbHNL with 40 and 71 changes, respectively, to make the protein sequence in the region surrounding the substrate-binding site identical to that in esterase SABP2. The substrate-binding site of SABP2 is defined as the region surrounding a bound product molecule, salicylic acid, in the x-ray structure of SABP2 (PDB entry 1y7i).

Variant HNL40 introduces 40 changes to HbHNL to increase its sequence similarity to SABP2. Since there are 146 differences between the sequences of HbHNL and SABP2, these 40 changes eliminate approximately one quarter (27%) of the differences, raising the overall sequence identity with SABP2 from 44% for HbHNL to 59% for HNL40. These changes are concentrated to the substrate-binding region and are hypothesized to make the catalytic atom positions better mimic those in SABP2.

The substrate binding region of SABP2 contains 59 residues with at least one atom within 10 Å of the bound salicylic acid. Of these, 27 residues are identical in SABP2 and *Hb*HNL, while 32 differ. To fully match the sequence of the substrate-binding region, HNL40 incorporates substitutions for all 32 differing residues. These 32 substitutions are as follows (*Hb*HNL numbering): I9V, T11G, I12A, I18S, E79H, C81L, L84M, H103L, N104A, S105A, V106F, V118L, L121Y, M122N, F125T, D127N, Y133F, L146M, K147F, L148F, L152F, N156K, T173V, G176S, Q180M, E208K, I209G, F210I, L211P, K236M, L237A, and Q238M.

HNL40 contains eight additional changes in the lid domain beyond this 10-Å region. There is an insertion of two residues, AE, after residue 126 and six additional substitutions (HbHNL numbering): K129L, F150P, T151K, R154A, E155H, and T159Q. Thus, HNL40 contains a total of 40 amino acid changes relative to *Hb*HNL: 38 amino acid substitutions and two inserted amino acids. In the HNL40 and HNL71 numbering, the numbering of the matching residues at positions 127 and above is two units higher than in *Hb*HNL due to the two-amino-acid-residue insertion after position 126.

Variant HNL71 contains 31 additional substitutions beyond those in HNL40 for a total of 71, which corresponds to removing approximately one half (49%) of the differences between the proteins. These 31 additional substitutions increase the sequence identity to SABP2 to 71% for HNL71 and expand the region of identical protein sequence with SABP2 to 14 Å from salicylic acid bound in the substrate-binding site of SABP2. There are 130 residues in SABP2 with at least one atom within 14 Å of the salicylic acid bound to the substrate-binding site. Fifty-nine of these are within 10 Å, so there are 71 more residues to consider in the region from 10 to 14 Å. Thirty-two of these 71 residues are identical between SABP2 and *Hb*HNL and 39 residues differ. Eight of these changes in the lid region were already included in HNL40 above, so only 31 additional substitutions are required to make the sequence identical to that in SABP2 within 14 Å of the substrate-binding site. Those additional 31 substitutions are (*Hb*HNL numbering): A16G, I51L, S53T, F54L, S58T, I86L, A87G, L107M, P114S, Y116F, D119E, K120Q, V124R, T132Q, F134L, G145S, E165D, Y166L, K170S, K175P, N181E, V202I, W203V, T204C, D205T, Q206E, P212E, L216R, G233A, T240C, I245L.

The x-ray structures of HNL40 and HNL71 reveal the degree to which these substitutions increased the similarity of the catalytic atom positions to those in SABP2. The 40 changes in HNL40 make these positions similar, but not identical; the additional 31 substitutions in HNL71 make the catalytic atom positions identical to those in SABP2. The tunnels in HNL40 and HNL71 connect the substrate-binding site, which contains residues identical to those in SABP2, and the enzyme surface, which contains residues differing from those in SABP2. The pattern of tunnels in HNL40 and HNL71 is intermediate between that in *Hb*HNL and SABP2.

## 2. Materials and Methods

### 2.1. Protein production and purification

The HNL40 and HNL71 genes containing a His-tag (N-terminal and C-terminal respectively) were codon-optimized, synthesized, and cloned into pET21a(+) vectors by Twist Biosciences (Table 1). The gene for HNL40 encodes 293 amino acids (259 amino acids plus 33 amino acids to encode an N-terminal linker and His6 tag), while the gene for HNL71 encodes 270 amino acids (259 amino acids plus 11 amino acids to encode a C-terminal linker and His6 tag). Table S1 in the Supporting Information contains the full sequences of the expression plasmids.

**Table 1.**
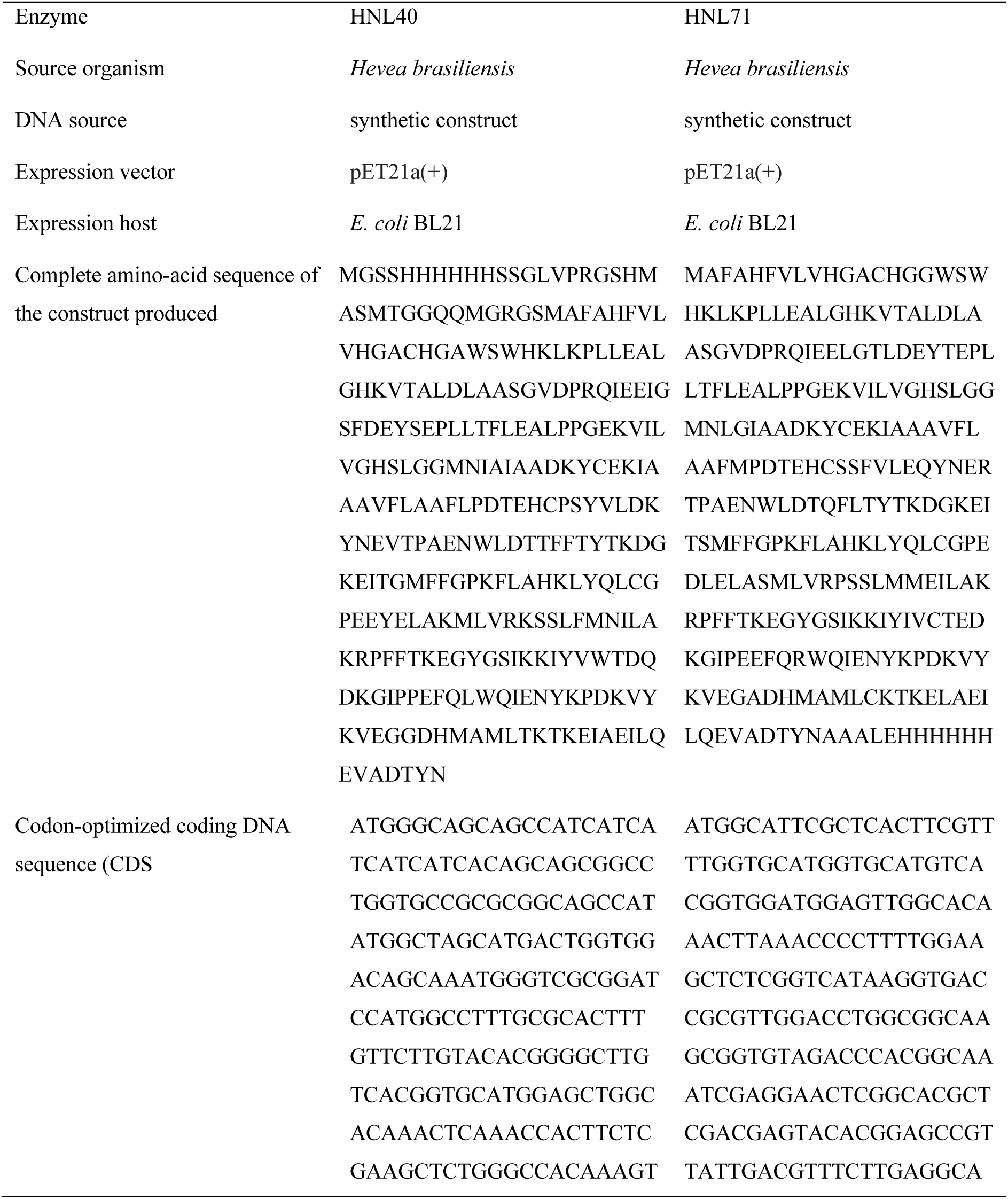
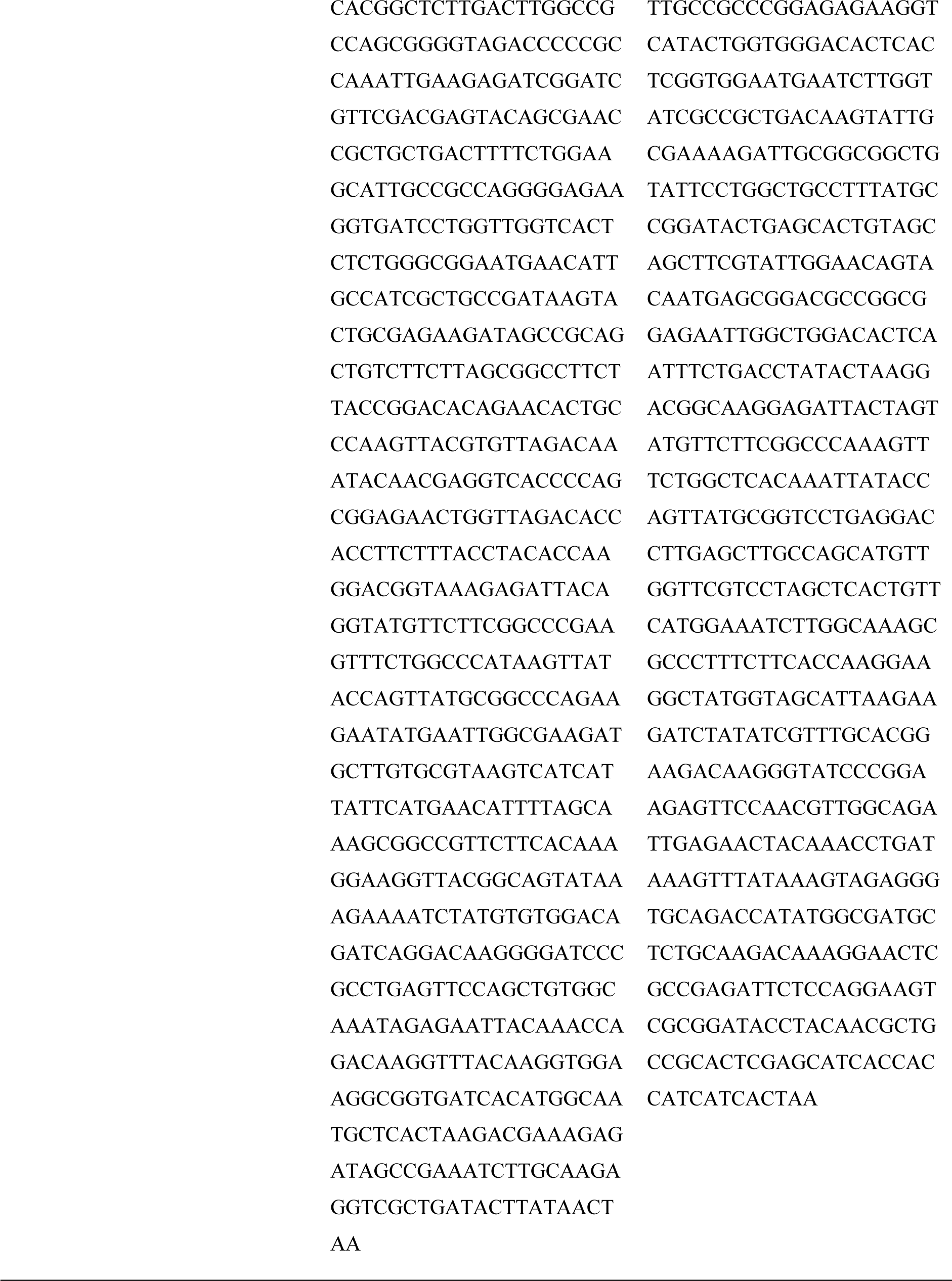
Protein-production information.

Lysogeny broth (LB) media (5 ml) containing carbenicillin (100 μg ml^−1^) was inoculated with a single bacterial colony picked from an agar plate and incubated in an orbital shaker at 37 °C and 240 rpm for 15 h to create a seed culture. A 1-liter baffled flask containing terrific broth-carbenicillin media (250 ml) was inoculated with 2.5 ml of seed culture. This pre-induction culture was incubated at 37 °C and 240 rpm for 3–4 h until the absorbance at 600 nm reached 0.4-1.0. The flask was then cooled on ice for 30 min. Isopropyl β-D-1-thiogalactopyranoside (0.75–1.0 m*M* final concentration) was added to induce protein expression, and cultivation was continued for 20-24 h at 16 °C. The cells were harvested by centrifugation (7000 rcf, 15 min at 4 °C), resuspended in NiNTA loading buffer (10 m*M* imidazole, 50 m*M* Tris pH 8.0, 500 m*M* NaCl, 4 ml g^−1^ of wet cells), and either directly sonicated or frozen for storage and later purification. Cells were flash-frozen in liquid nitrogen or a dry ice-ethanol bath and stored at −80 °C. Frozen cells were thawed at room temperature or in a room temperature water bath, and fresh/thawed cells were disrupted by sonication (400 W, 40% amplitude for 3 min). The cell lysate was centrifuged to pellet the cell debris (4 °C, 20,000 rcf for 20 min), which was discarded.

The crude lysate was purified via nickel affinity chromatography. The supernatant from above was mixed with 1-2.5 ml of NiNTA resin (pre-equilibrated with 10 ml of NiNTA loading buffer) and incubated for 45 min at 4 °C with rotation (10 rpm). The resin/supernatant mixture was loaded onto a 25-ml gravity flow column (Bio-Rad) and the resin was washed with 10 column volumes each of buffers A and B for a total of 20 column volumes (50 m*M* Tris pH 8.0, 500 m*M* NaCl, buffer A contained 25 m*M* imidazole and buffer B contained 50 m*M* imidazole). The His-tagged protein was eluted with 10 column volumes of elution buffer (125 m*M* imidazole, 50 m*M* Tris pH 8.0, 500 m*M* NaCl) and collected in 1-ml fractions. The protein concentration of each elution fraction was determined from its absorbance at 280 nm measured using a Nanodrop 2000 (Thermo Scientific) and the calculated extinction coefficient for the protein (https://web.expasy.org/protparam/). Protein gels were used to check for the presence and purity of protein and run using sodium dodecyl sulfate-polyacrylamide gradient gels (NuPage 4−12% Bis-Tris gel from Invitrogen) using the Precision Plus Dual Color protein standard (BioRad, 5 μl per lane), run for 50-60 min at 120V, stained with SimplyBlue Safe Stain (Thermo Fisher Scientific), and destained 2x with Milli-Q UltraPure water. SDS-PAGE indicated a molecular weight of ∼31 kDa in agreement with the predicted weights of 32.8 kDa for the HNL40 construct and 30.4 kDa for the HNL71 construct. Only the most concentrated elution fractions, typically fractions 2-10, were pooled to reduce the presence of contaminating proteins and then sterile-filtered. The imidazole-containing elution buffer was exchanged by the addition of BES buffer (5 m*M N*, *N*-bis(2-hydroxyethyl)-2-aminoethanesulfonic acid (BES), pH 7.2, 14 ml) followed by ultrafiltration (Amicon 15-ml ultrafiltration centrifuge filter, 10 kDa cutoff) to reduce the volume to ∼1 ml. This addition of buffer and filtration was repeated four times. The last ultrafiltration spin was extended by 10 min to reduce the volume to ∼250 μl. The final protein concentration was determined via spectrophotometric measurements at 280 nm, measured in duplicate, and averaged. A 250-ml culture typically yielded 2-5 mg of protein.

Samples used for crystallization were further purified via size exclusion chromatography. The purification column (Cytiva HiLoad™ 16/600 Superdex™ 200 pg, 120-ml capacity) was equilibrated with 1.10 column volumes of running buffer (5 m*M* BES, pH 7.2) prior to sample injection. The sample was injected and eluted at a flow rate of 1 ml min^−1^ for 1.10 column volumes of running buffer. Three-milliliter fractions were collected in 15-ml conical tubes when the A_280_ signal intensity exceeded 15 mAU (absorbance units). HNL71 eluted at 65 - 75 min and signal intensity peaked at ∼70 min while HNL40 peaked at ∼64 minutes (Figure S1). Elution fractions 2-6 (HNL71) and 1-5 (HNL40) were evaluated for purity, sterile-filtered, pooled, and concentrated as above. The final, purified protein concentrations were 12 mg ml^−1^ (HNL71) and 8 mg ml^−1^ (HNL40).

### 2.2. Crystallization

Crystallographic screening was done using the Crystal Phoenix Dispenser (Art Robbins Instruments/Hudson Robotics). Sitting drop trays and the CrystalMation Intelli-Plate low-profile 96 Well plates from Hampton Research were used for the crystallization setup. All the setup and washing procedures were done through the Phoenix software.

For HNL40, each crystallization drop consisted of 0.1 μl protein sample (8 mg ml^−1^ protein) mixed with 0.1 μl well solution (Table 2). A total of 960 conditions were screened. Crystals formed within 1 day from the Index HT screen from Hampton Research Inc under the condition of 0.1 *M* 4-(2-hydroxyethyl)-1-piperazineethanesulfonic acid (HEPES), pH 7.5, 0.2 *M* L-proline, and 10% (*w/v)* polyethylene glycol 3,350 and grew to the full size of 160 µm x 50 µm x 10 µm in 3 days. The cryoprotectant solution for the HNL40 crystals comprised 0.1 *M* HEPES, pH 7.5, 0.2 *M* L-proline, 10% (*w/v)* polyethylene glycol 3,350, and 25% (*v/v*) ethylene glycol. This initial robotic screening produced crystals sufficient for model refinement.

**Table 2.** Crystallization information.

| Enzyme | HNL40 | HNL71 |
| --- | --- | --- |
| Method | Sitting Drop | Sitting Drop |
| Plate type | CrystalMation Intelli-Plate low-profile 96 Well, Hampton Research | CrystalMation Intelli-Plate low-profile 96 Well, Hampton Research |
| Temperature (K) | 298 | 298 |
| Protein concentration (mg ml<sup>-1</sup>) | 8 | 12 |
| Buffer composition of protein solution | 5 mM BES buffer, pH 7.2 | 5 mM BES buffer, pH 7.2 |
| Composition of reservoir solution | 0.1 M 4-(2-hydroxyethyl)-1-piperazine-ethanesulfonic acid (HEPES), pH 7.5, 0.2 M L-proline, 10% (w/v) polyethylene glycol 3,350 | 0.1 M Tris pH 8.5, 0.2 M magnesium chloride hexahydrate, 25% (w/v) polyethylene glycol 3,350 |
| Volume and ratio of drop (μl) | 0.2, 1:1 | 0.2, 1:1 |
| Volume of reservoir (μl) | 50 | 50 |

For HNL71, each crystallization drop contained 0.1 μl protein sample (12 mg ml^−1^ protein) mixed with 0.1 μl well solution (Table 2). A total of 960 conditions were screened. Crystals formed within 1 day from the Index HT screen from Hampton Research Inc under the condition of 0.1 *M* Tris pH 8.5, 0.2 *M* magnesium chloride hexahydrate, and 25% (*w/v)* polyethylene glycol 3,350 and grew to the full size of 50 µm x 80 µm x 30 µm in 3 days. The cryoprotectant solution for the HNL71 crystals comprised 0.1 *M* Tris pH 8.5, 0.2 *M* magnesium chloride hexahydrate, 25% (*w/v)* polyethylene glycol 3,350, and 25% (*v/v*) ethylene glycol. This initial robotic screening produced crystals sufficient for model refinement.

### 2.3. Data collection, processing, and structure refinement

The x-ray diffraction data for HNL40 were collected at the Advanced Photon Source (Lemont, Il) on beamline 24-ID-C using a wavelength of 0.979 Å, Table 3. Crystals were preserved by flash-cooling in liquid nitrogen with a cryoprotectant solution comprised of 0.1 *M* HEPES, pH 7.5, 0.2 *M* L-proline, 10% (*w/v)* polyethylene glycol 3,350, and 25% (*v/v*) ethylene glycol. Data were collected at 100 K with a DECTRIS EIGER2 S 16M detector, with an exposure time of 0.2 seconds per frame, a crystal-to-detector distance of 180 mm, and an oscillation angle of 0.2° per frame. The crystal belonged to the space group *P*2_1_2_1_2_1_ with unit cell dimensions a = 76.527 Å, b = 81.884 Å, and c = 90.929 Å. The dataset was 98.04% complete to a resolution of 1.99 Å, with an overall R_merge_ of 0.066 and a multiplicity of 3.70.

**Table 3.**
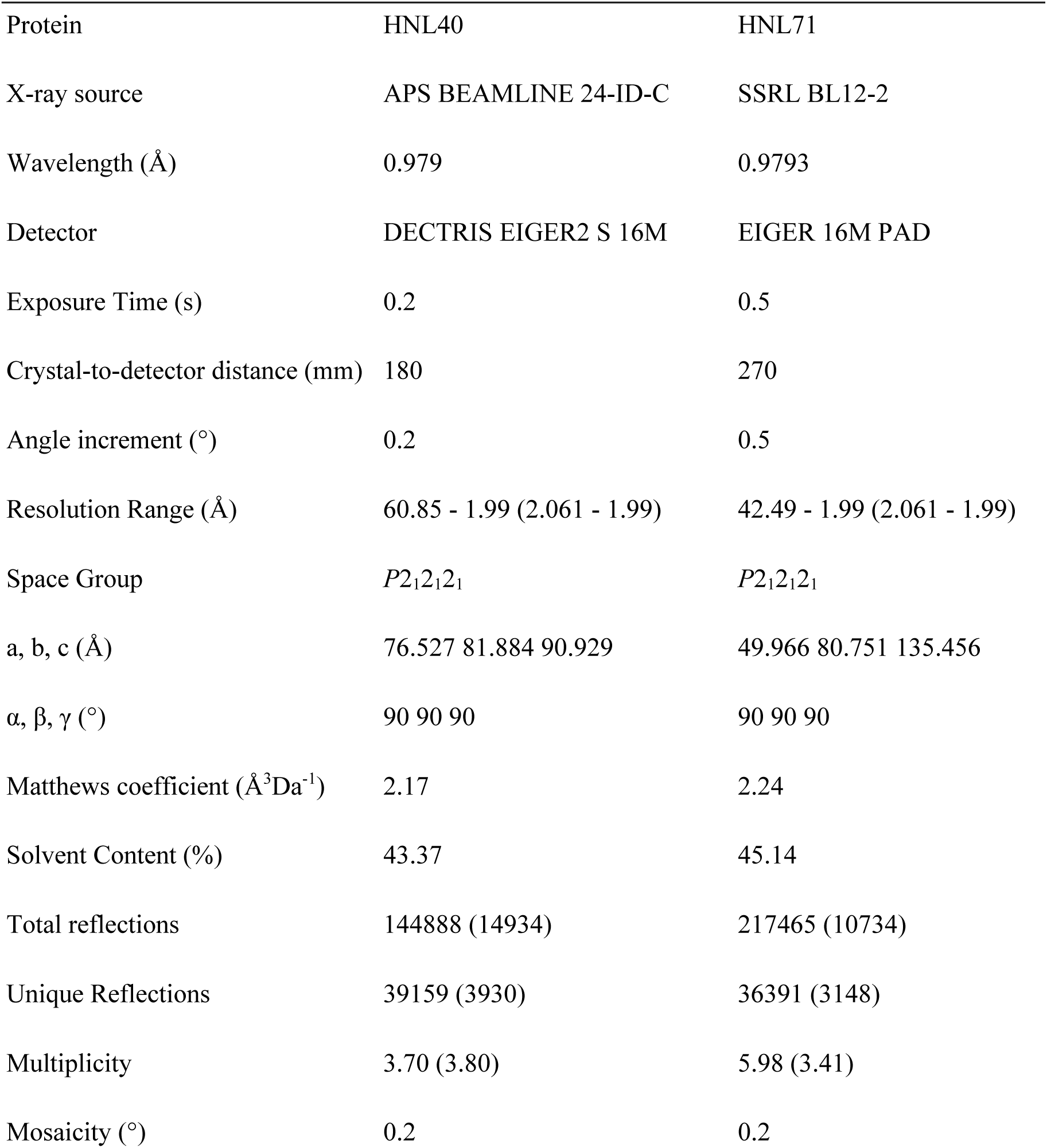
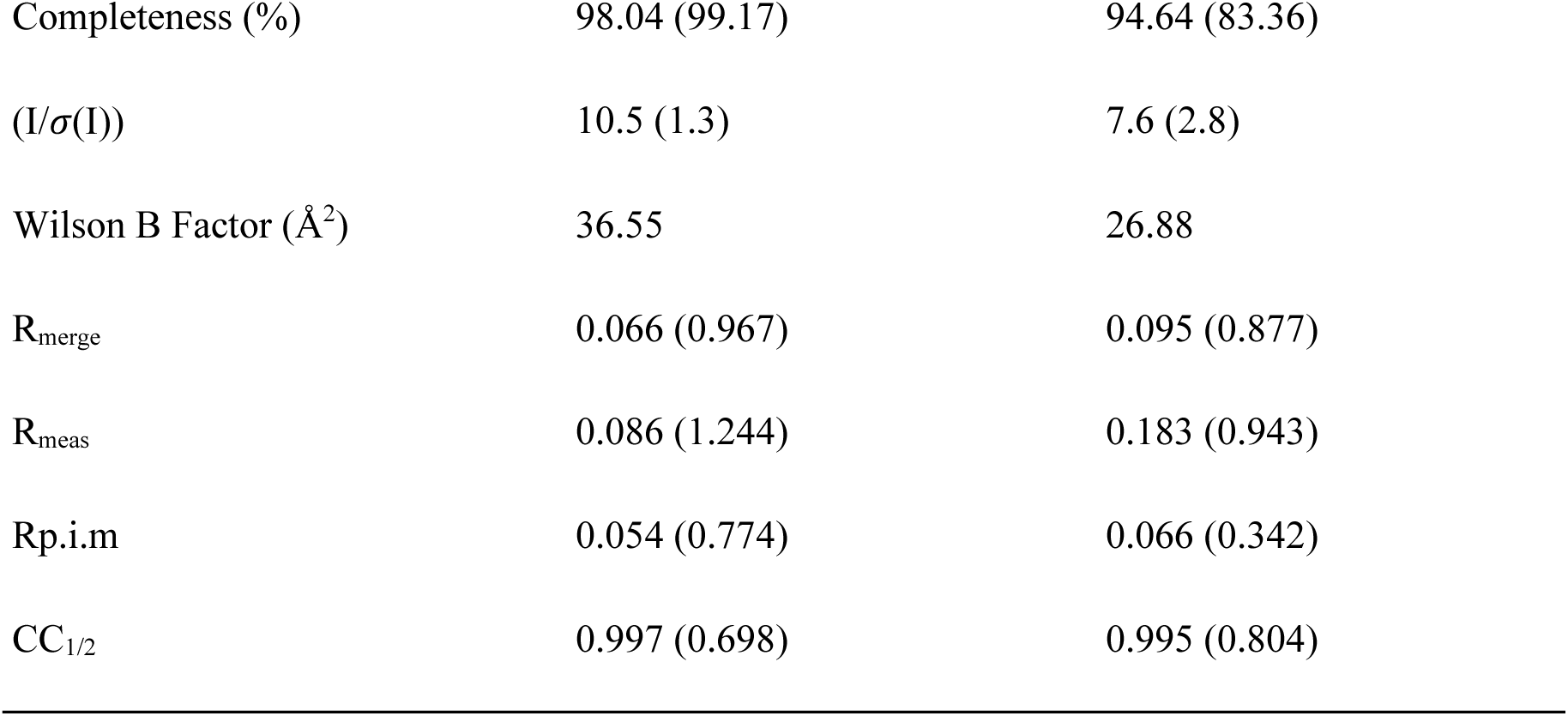
Data collection and processing.

The x-ray diffraction data for HNL71 were collected at the Stanford Synchrotron Radiation Lightsource (SSRL) on beamline BL12-2 using a wavelength of 0.9793 Å. The crystals were preserved by flash-cooling in liquid nitrogen using a cryoprotectant solution composed of 0.1 *M* Tris pH 8.5, 0.2 *M* magnesium chloride hexahydrate, 25% (*w/v*) polyethylene glycol 3,350, and 25% (*v/v*) ethylene glycol. Data were collected at 100 K with an EIGER 16M PAD detector, with an exposure time of 0.5 seconds per frame, a crystal-to-detector distance of 270 mm, and an oscillation angle of 0.5° per frame. The crystal that generated the dataset belonged to the space group *P*2_1_2_1_2_1_ with unit cell dimensions a = 49.966 Å, b = 80.751 Å, and c = 135.456 Å. The dataset was 94.64% complete to a resolution of 1.99 Å, with an overall R_merge_ of 0.095 and a multiplicity of 5.98. The final data collection and processing statistics are summarized in Table 3.

The data for both crystals were processed using the HKL2000 (Otwinowski & Minor, 1997). The initial structure of HNL40 was obtained via molecular replacement using the structure of hydroxynitrile lyase from rubber tree with seven mutations (PDB entry 8euo) as the search model with an inputted sequence identity of 84.5%. The initial structure of HNL71 was obtained via molecular replacement using the predicted structure of HNL71 from AlphaFold2 (Jumper *et al*., 2021), which was used as the search model with an inputted sequence identity of 100%. This 100% value ignores the C-terminal His6-tag in HNL71, which was not resolved in the electron density. Molecular replacement was performed with Phaser (McCoy *et al*., 2007) in the Phenix software suite (Liebschner *et al*., 2019). Subsequent rounds of refinement were carried out using phenix.refine and manual adjustment of the initial model using Coot (Emsley *et al*., 2010). The software PDB-REDO (Joosten *et al*., 2014) was used at an intermediate refinement stage (round 66) for HNL71 and followed by additional manual refinement to ensure a chemically reasonable structure. PDB-REDO was not used during the refinement of the HNL40 structure.

Tables 3 and 4 show the detailed collection, processing, and refinement statistics for both proteins. The figures of the protein structure in this paper were created using PyMOL v.2.5.4 or v.3.03 (Schrödinger, LLC., 2024; https://pymolwiki.org). The final structure models were deposited in the Research Collaboratory for Structural Bioinformatics Protein Data Bank as PDB entry 8sni for HNL40 and PDB entry 9clr for HNL71.

## 3. Results and Discussion

### 3.1. Crystallization and structure determination

Molecular replacement using the x-ray structure of a homolog (84.5% sequence identity, 8euo) generated an initial structure for HNL40 and refinement based on 39,116 reflections with 2,007 reflections (5%) set aside for R_free_, yielded a final structure with an R_work_ of 0.1991 and an R_free_ of 0.2233. The asymmetric unit contained two protein monomers as did the structure of the parent protein *Hb*HNL (PDB ID 1yb6, Wagner *et al*., 1996) and the target protein SABP2 (PDB ID 1y7i, Forouhar *et al*., 2005). The dimer consists of 517 protein residues, 68 solvent molecules, and a total of 4,013 non-hydrogen atoms. The average B-factor for the final model was 38.61 Å². Structural validation using MolProbity showed 97.27% of the residues in the favored regions and 2.73% in the allowed regions of the Ramachandran plot, with no outliers (Table 4).

**Table 4.**
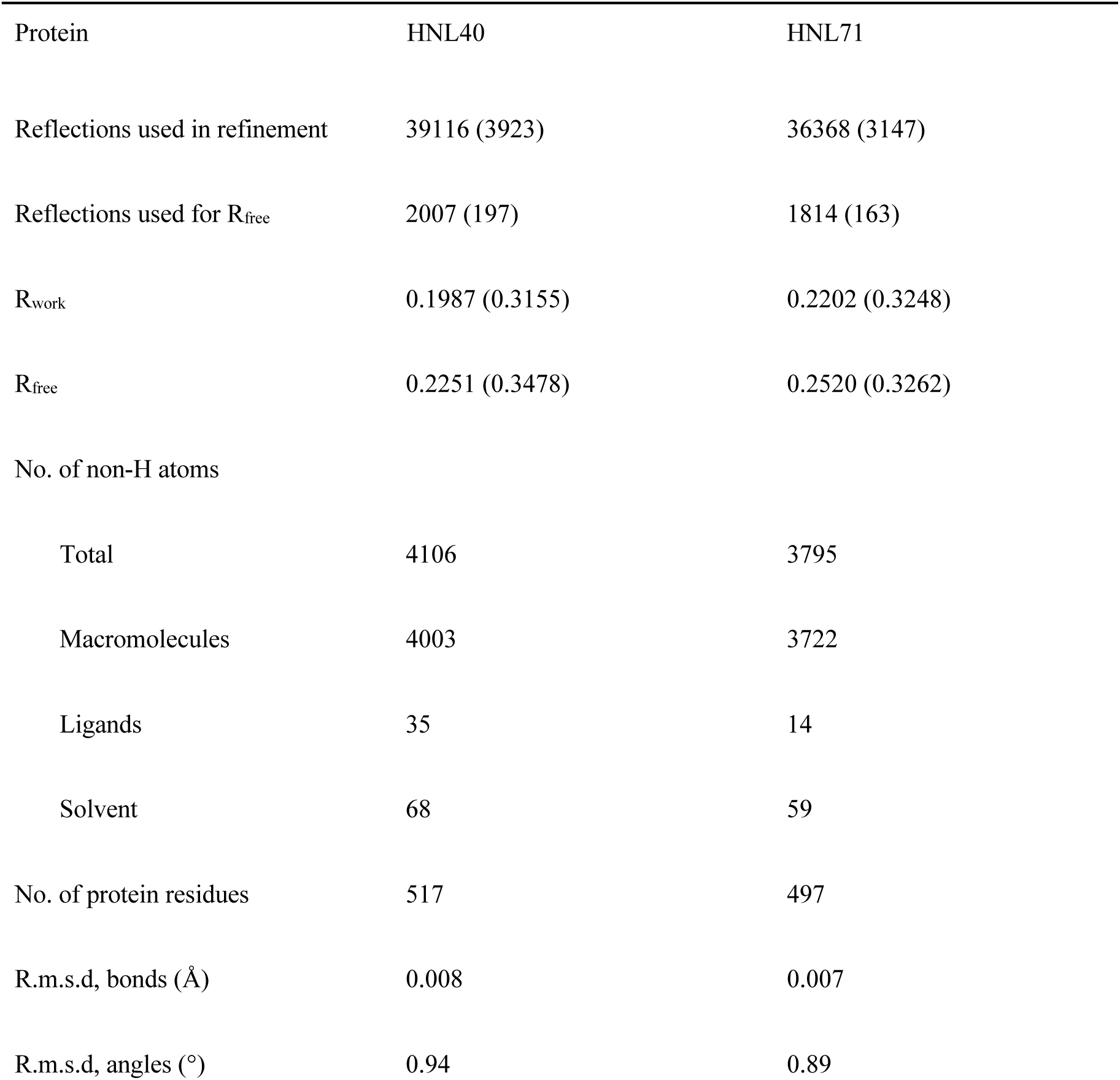
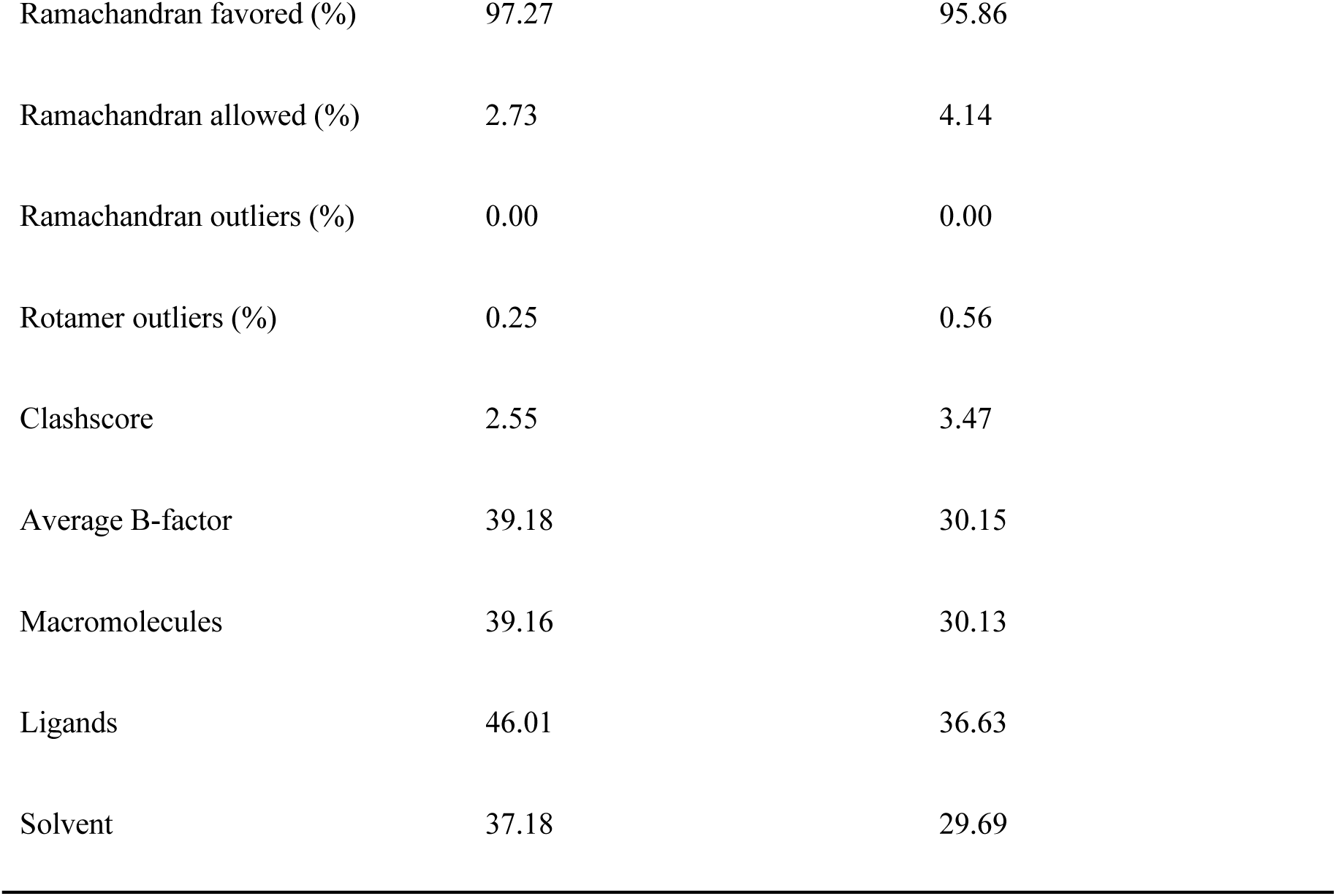
Structure refinement information for HNL40 and HNL71 Values in parentheses are for the highest-resolution shell.

Molecular replacement using an AlphaFold2 (Jumper *et al*., 2021) model of HNL71 generated an initial structure of HNL71 with an LLG of 3623.325 and a TFZ of 59. Refinement of the structure over 98 rounds was based on 36,368 reflections with 1,814 reflections (5%) set aside for R_free_. PDB-REDO (Joosten *et al*., 2014) was used at an intermediate refinement stage and improved the fit from R_work_ of 0.2293 and R_free_ of 0.2615 to a R_work_ of 0.2281 and a R_free_ of 0.2590. The final structure had an R_work_ of 0.2202 and an R_free_ of 0.2520. As with HNL40, there were two protein monomers in the asymmetric unit of HNL71. The dimer consists of 497 protein residues, 59 solvent molecules, and a total of 3,795 non-hydrogen atoms. The average B-factor for the final model was 30.15 Å². Structural validation using MolProbity showed 95.86% of the residues in favored regions and 4.14% in the allowed regions of the Ramachandran plot, with no outliers (Table 4).

As it is common in x-ray structures, the N-terminal His6 tag and linker in HNL40 and the C-terminal His6 tag and linker in HNL71 were not seen in the electron density. In HNL71, the data also lacked sufficient electron density to define nine residues in two surface regions: a two-residue segment Gly142Lys143 in the cap domain, and a seven-residue segment Met182Glu183Ile184Leu185Ala186Lys187Arg188 in the catalytic domain.

The structures of esterase SABP2, hydroxynitrile lyase *Hb*HNL and variants HNL40 and 71 are similar and consist of two domains, as shown in Fig. 3. According to the *Hb*HNL numbering, the catalytic domain (1-107, 180-265) contains the catalytic triad (Ser80-Asp207-His237) and adopts the α/β-hydrolase fold, while the lid domain (108-179) forms part of the substrate-binding region. Fig. 3 also shows, superimposed on the structure of HNL40, the locations of the substitutions and insertions that created HNL40 and HNL71 from *Hb*HNL. The 38 substitutions in HNL40 surround the substrate binding site and the two inserted amino acids lie on the surface in the lid domain. The additional substitutions in HNL71 further extend the region of sequence identity between SABP2 and the variant protein.

**Figure 3.**
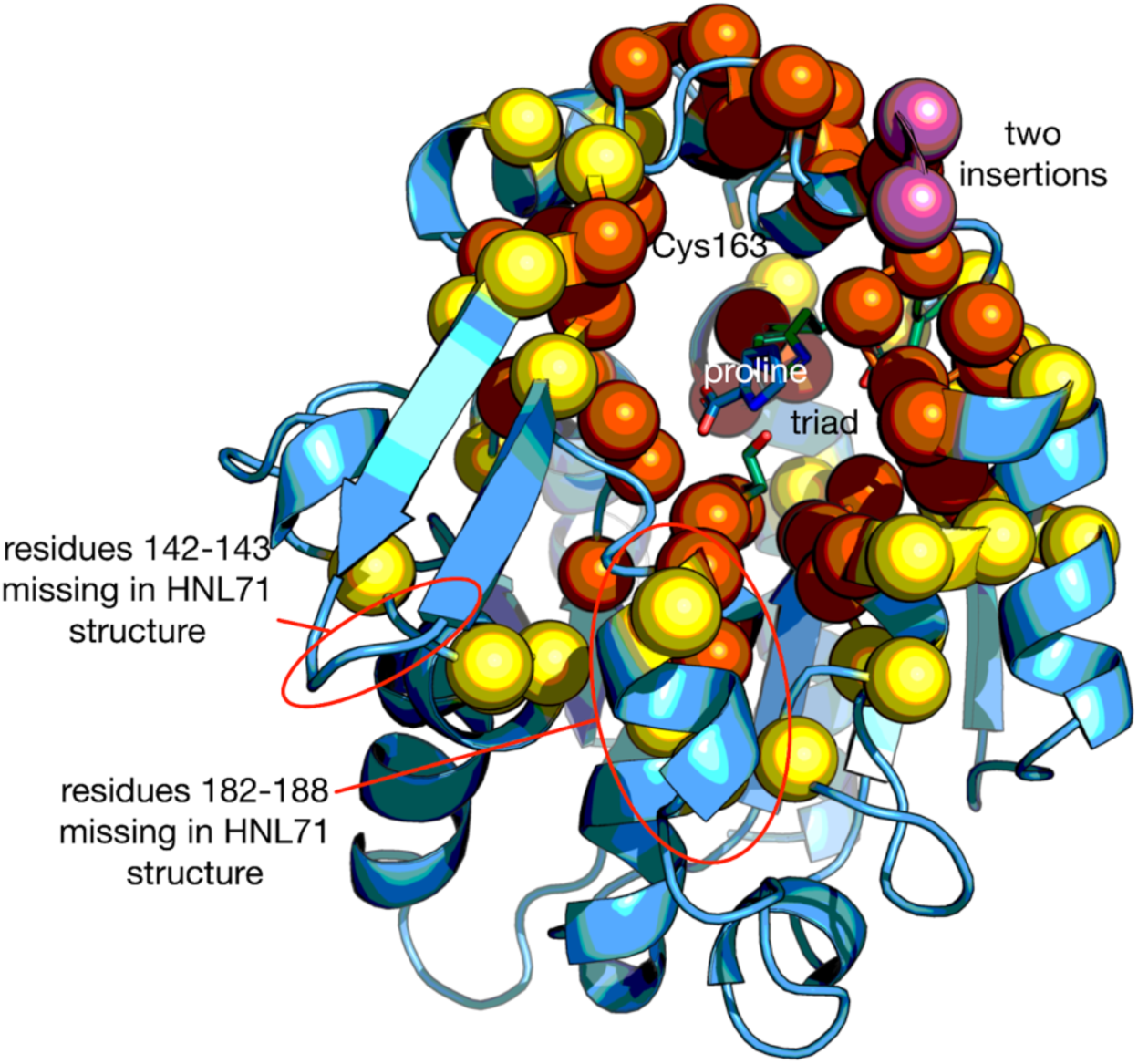
Ribbon diagram of the structure of chain A of HNL40 (PDB entry 8sni) showing the locations of changes introduced to *Hb*HNL to create HNL40 and HNL71. The lid domain (108-181) lies at the top and the catalytic domain (1-107, 182-267) lies at the lower part in this view. The catalytic triad Ser80-His237-Asp209 (side chains shown as green sticks) lies in the catalytic domain at the interface between the two domains. A bound L-proline (blue sticks) lies in the substrate binding region and mimics the product carboxylate of an ester hydrolysis. Both HNL40 and HNL71 include two insertions (rose spheres) after residue 126 of *Hb*HNL and the 38 substitutions marked as dark orange spheres. These changes in HNL40 make the protein sequence of its region within 10 Å of the substrate-binding site identical to that in SABP2. HNL71 contains 31 additional substitutions (yellow spheres) to make the protein sequence of the region within 14 Å identical to that in SABP2. In the HNL71 structure, the electron density was insufficient to define the positions of nine surface-exposed amino acid residue: 142-3 (lid domain) and 182-8 (catalytic domain). These regions were defined in the HNL40 structure shown and are circled in red. In HNL71, the side chain of Cys163 (sticks) is oxidized to the sulfenic acid.

### 3.2. Catalytic residue positions

The catalytic residue positions were compared in two ways (Table 5). The Cɑ atom positions compared the main chain positions, while the catalytic atom positions compared the side chain positions for the triad (Ser Oɣ, His Nε2, Asp Oδ2) and the main chain nitrogen positions for the two oxyanion hole residues. Three structures of SABP2 showed an average deviation of 0.3±0.1 Å in the Cɑ atom positions and 0.5±0.6 Å for the catalytic atom positions. Deviations of the atom positions by these amounts or less will be considered to have positions indistinguishable from those in SABP2.

**Table 5.**
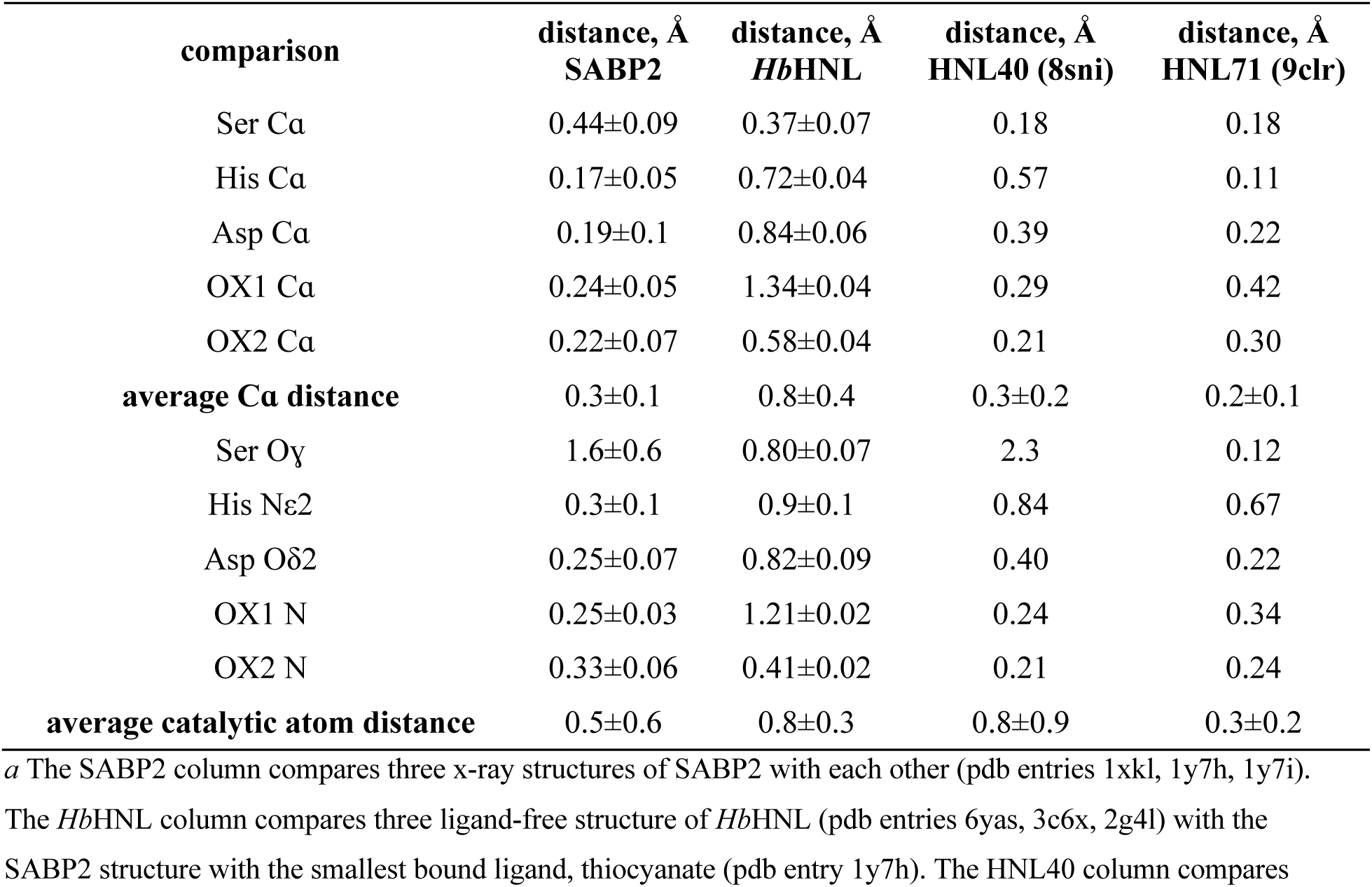
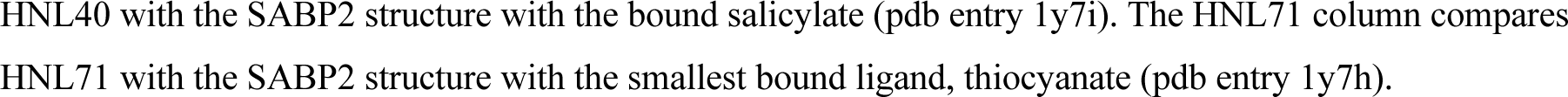
Deviation of Cɑ atom and catalytic atompositions of the oxyanion hole and catalytic triad residues between three structures of SABP2 and between SABP2 and *Hb*HNL, HNL40 and HNL71.

| comparison | distance, Å<br>SABP2 | distance, Å<br>*HbHNL* | distance, Å<br>HNL40 (8sni) | distance, Å<br>HNL71 (9clr) |
| --- | --- | --- | --- | --- |
| Ser Ca | 0.44±0.09 | 0.37±0.07 | 0.18 | 0.18 |
| His Ca | 0.17±0.05 | 0.72±0.04 | 0.57 | 0.11 |
| Asp Ca | 0.19±0.1 | 0.84±0.06 | 0.39 | 0.22 |
| OX1 Ca | 0.24±0.05 | 1.34±0.04 | 0.29 | 0.42 |
| OX2 Ca | 0.22±0.07 | 0.58±0.04 | 0.21 | 0.30 |
| **average Ca distance** | 0.3±0.1 | 0.8±0.4 | 0.3±0.2 | 0.2±0.1 |
| Ser Oγ | 1.6±0.6 | 0.80±0.07 | 2.3 | 0.12 |
| His Nε2 | 0.3±0.1 | 0.9±0.1 | 0.84 | 0.67 |
| Asp Oδ2 | 0.25±0.07 | 0.82±0.09 | 0.40 | 0.22 |
| OX1 N | 0.25±0.03 | 1.21±0.02 | 0.24 | 0.34 |
| OX2 N | 0.33±0.06 | 0.41±0.02 | 0.21 | 0.24 |
| **average catalytic atom distance** | 0.5±0.6 | 0.8±0.3 | 0.8±0.9 | 0.3±0.2 |

<sup>a</sup> The SABP2 column compares three x-ray structures of SABP2 with each other (pdb entries 1xkl, 1y7h, 1y7i).
The *HbHNL* column compares three ligand-free structure of *HbHNL* (pdb entries 6yas, 3c6x, 2g4l) with the SABP2 structure with the smallest bound ligand, thiocyanate (pdb entry 1y7h). The HNL40 column compares
HNL40 with the SABP2 structure with the bound salicylate (pdb entry 1y7i). The HNL71 column compares HNL71 with the SABP2 structure with the smallest bound ligand, thiocyanate (pdb entry 1y7h).

The catalytic residue positions in *Hb*HNL (three ligand-free structures) differ from those in SABP2 by 0.8±0.4 Å for the Cɑ atom positions and 0.8±0.3 Å for the catalytic atom positions, Table 5 and Fig. 2 above. The largest deviation is in the position of Cɑ and N of the OX1 residue, 1.2-1.3 Å. The two proteins therefore subtly differ in the position of their catalytic machinery.

For HNL40, the average Cɑ atom positions, but not the catalytic atoms positions, match those in the three SABP2 structures. The average Cɑ atom distance, 0.3±0.2 Å, is similar to that for comparison of the three SABP2 structures to each other. However, the average catalytic atom distances of 0.8±0.9 Å is larger than the comparison of SABP2 structures to each other, 0.5±0.6 Å. The largest deviations are 2.3 Å for Ser Oɣ and 0.84 Å for His Nε2. Thus, the forty changes introduced to create HNL40 are insufficient to create identical catalytic atom positions to those in SABP2.

For HNL71, both the average Cɑ atom positions and the catalytic atom positions match those in the three SABP2 structures. The average distances between HNL71 and SABP2 are smaller than those for a comparison of three SABP2 structures to each other. However, comparison of the individual distances suggest that the His Nε2 and OX1 N positions in HNL71 still differ from those in SABP2. Thus, the seventy-one changes introduced to create HNL71 create nearly, but not completely, identical catalytic atom positions.

### 3.3. L-Proline bound in the active site of HNL40

The structure of HNL40 contains an L-proline bound in the substrate-binding pocket, Fig. 3 above. Fig. S2 shows the electron density map that defines the bound L-proline. The L-proline originated from the crystallization solution, which contained 0.2 M L-proline, and mimics a bound carboxylate product of an esterase reaction. At the near neutral pH of the crystallization solution, proline is expected to be zwitterionic with a negatively charged carboxylate (p*K_a_* 1.99) and positively charged ammonium group (p*K_a_* 10.96).

In the SABP2 structure containing a bound salicylate, the salicylate carboxylate accepts hydrogen bonds from the catalytic atoms, Fig. 4, Table S2. The catalytic serine Oɣ donates hydrogen bonds to both carboxylate oxygens, the catalytic histidine Nε2 donates a hydrogen bond to one carboxylate oxygen, and the two oxyanion hole N-H’s donate hydrogen bonds to the other carboxylate oxygen.

**Figure 4.**
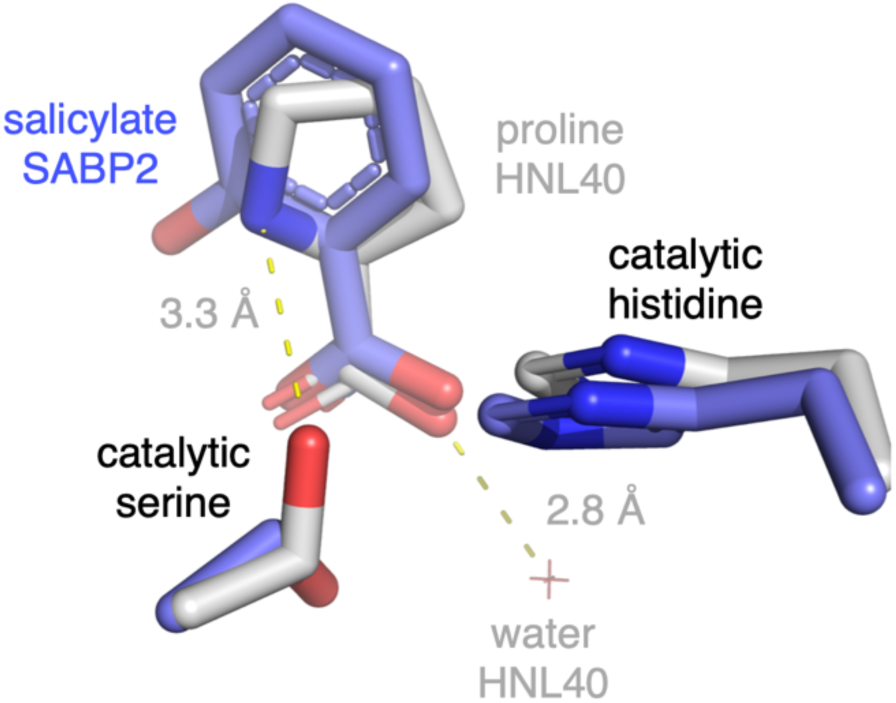
Overlay of the structure of SABP2 with a bound product salicylate (PDB entry 1y7i, blue carbons) onto the structure of HNL40 with a bound proline (PDB entry 8sni, white carbons). The carboxylate moieties interact with the corresponding protein active sites slightly differently. The catalytic serine donates hydrogen bonds (not shown) to both salicylate carboxylate oxygens (3.0 Å, 3.0 Å) in the SABP2 structure, but the altered catalytic serine side chain conformation in the HNL40 structure makes the angle too acute for the serine to donate a hydrogen bond to the proline carboxylate even though the O-O distance is still favorable (3.0 Å, 3.0 Å). Instead, the one proline carboxylate oxygen accepts a hydrogen bond from a nearby water molecule (2.8 Å) and the proline ammonium donates a weak hydrogen bond to the catalytic serine (3.3 Å).

In the HNL40 structure containing the bound proline the hydrogen bond pattern with the catalytic serine differs because the catalytic serine Oɣ adopts a different conformation. It points toward the ammonium atom of the proline (3.3 Å) and now lies above the plane defined by the O–C–O of the proline carboxylate. This orientation relative to the lone pairs on the oxygens is too acute for a hydrogen bond. Instead, a water molecule donates a hydrogen bond to one of the carboxylates (2.8 Å). These differences in hydrogen bonding pattern between SABP2 and HNL40 appear to be due to the ammonium ion in proline rather than due to differences in the active site structure. Table S2 lists the relevant distances between atoms.

### 3.4. Similarity of the 10-Å region surrounding the substrate-binding site to the corresponding region in SABP2

As expected, the main chain positions of the *Hb*HNL variants become more similar to those of SABP2 as the sequence identity of the *Hb*HNL variants to SABP2 increases (Table 6). First, three different SABP2 structures were aligned with each other, which yielded an average RMSD of 0.30±0.05 Å over 216-235 of the 256 Cα atoms. The Cɑ positions of SABP2 and *Hb*HNL differ by more than twice that value – 0.64 Å over 227 out of 253 Cɑ atoms. The differences between SABP2 and HNL40 are smaller, 0.51 Å, and those for HNL71 even smaller, 0.41 Å. At the same time, the main chain positions in HNL40 and HNL71 move away from their original positions in *Hb*HNL. The RMSD between *Hb*HNL and HNL40 is 0.40 Å and increases to 0.47 Å in HNL71. HNL40 and HNL71 differ from each other with an RMSD of 0.30 Å.

**Table 6.** RMSD values for the alignment of all 256 Cɑ atoms in chain A of SABP2, *Hb*HNL, HNL40 and HNL71.

|  | SABP2 (1Y7I) | *Hb*HNL (1YB6) | HNL40 (8SNI) | HNL71 (9CLR) |
| --- | --- | --- | --- | --- |
| **SABP2**<br>(1y7h, 1xkl) | 0.30±0.05 Å for 216-235 atoms of 256 Ca atoms<sup>a</sup> | 0.64 Å for 227 of 253 Ca atoms | 0.51 Å for 235 of 255 Ca atoms | 0.41 Å for 219 of 246 Ca atoms |
| ***Hb*HNL** |  | — | 0.40 Å for 214 of 256 Ca atoms | 0.47 Å for 212 of 247 Ca atoms |
| **HNL40** |  |  | — | 0.30 Å for 207 of 249 Ca atoms |

<sup>a</sup> Three alignments of x-ray crystal structures of SABP2 (PDB entries 1y7i, 1y7h, 1xkl) with one another.

The largest differences in the Cα positions between *Hb*HNL and SABP2 occur in four regions (Fig. 5). The largest difference, 11.1 Å, occurs within a lid domain loop at SABP2 Pro142, which corresponds to Asp139 in *Hb*HNL. This difference and the differences for residues 140-145 in this loop are so large that they are divided by 4 to create the images in Fig. 5 so that the other differences in the Cα positions would also be clearly visible. The second region showing large differences between the Cα positions occurs near the two-amino-acid insertion (Ala130-Glu131 in SABP2). The Cα positions at the amino acid before the insertion, Pro129 in SABP, Pro126 in *Hb*HNL, differ by 5.6 Å. Fig. 5 shows the inserted amino acids as a dotted line because there is no corresponding amino acid in *Hb*HNL making ΔCα undefined. The third region showing large differences between the Cα positions is surface loop 188-195 in SABP (185-191 in *Hb*HNL). The loop conformation differs with the largest Cα difference (∼3 Å) occurring at SABP2 Ala189-Lys190-Tyr191. Finally, the Cα positions of the N- and C-termini of the two proteins differ by 6 Å at Glu3 of SABP2 and by 3 Å at Asn260 of SABP2.

**Figure 5.**
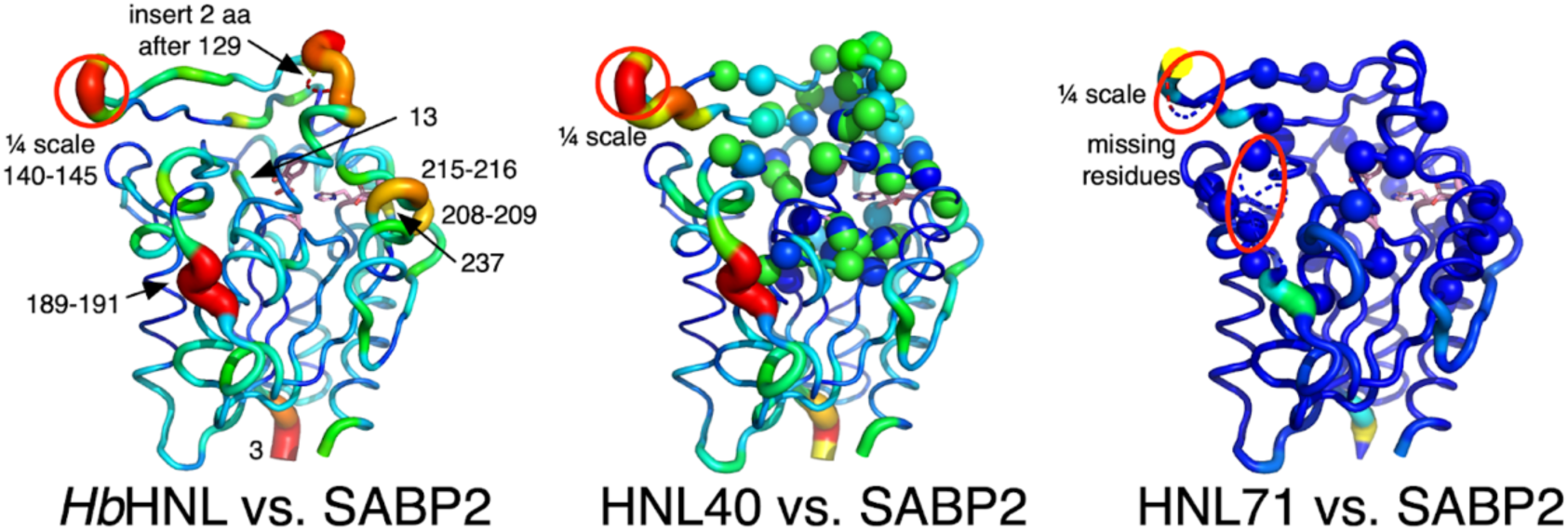
Displacement of Cα atoms (ΔCα) in *Hb*HNL (PDB entry 1yb6), HNL40 (PDB entry 8sni), and HNL71 (PDB entry 9clr) relative to wt SABP2. While the structures of SABP2 and *Hb*HNL differ significantly, the addition of 40 changes to *Hb*HNL to create HNL40 makes the region near the active site more similar to SABP2 and the 71 changes in HNL71 make this region identical to SABP2. All three images show the structure of SABP2 (PDB entry 1y7i), but the color and cartoon thickness indicate the displacement of the Cα atoms in the other structures. The triad and bound salicylic acid are shown as pink sticks. The representations are scaled so that a red, thick sausage indicates a displacement of ∼3 Å in all three structures, with the exception of residues 136-141 (circled) in the lid region of SABP2. The Cα positions in this region differ by up to 11 Å from that in *Hb*HNL, so this representation was scaled down for clarity. The dotted line near the top in the *Hb*HNL comparison indicates the two-amino-acid insertion in SABP2 that has no counterpart in *Hb*HNL. The two dotted line regions in the HNL71 comparison (marked by red ovals) were not observed in the x-ray structure suggesting disorder. The HNL40 comparison shows the Cα atoms as spheres at the locations of the 40 changes. The HNL71 comparison shows its additional 31 substitutions as spheres at the Cα positions.

Three other regions with smaller ΔCα differences between *Hb*HNL and SABP2 are noteworthy because they involve the catalytic residues. The Cα positions of oxyanion hole residue OX1 (Ala13 in SABP2; Ile12 in *Hb*HNL) differ by 1.3 Å. The Cα position of residue Asp237 in SABP2 near the catalytic His238 differs by 1.8 Å from the corresponding Asp234 in *Hb*HNL. The Cα position of residues 208-209 next to the catalytic Asp210 differ by 2.0 Å, as well as residues 215-216 differ by 2.2 and 2.0 Å, which are adjacent to 208-209.

Adding two insertions and 38 substitutions to create HNL40 significantly reduces the differences between it and SABP2 near the two-amino-acid insertion, and all regions involving the catalytic residues (Fig. 5). The differences at the lid domain loop at SABP2 Pro142, surface loop 188-195 in SABP2, and N- and C-termini remain largely unchanged. Adding an additional 31 substitutions to create HNL71 further reduces differences near the two-amino-acid insertion and all regions involving the catalytic residues (Fig. 5). The x-ray structure did not resolve part of the lid domain loop at SABP2 Pro142 and surface loop 188-195 in SABP2, suggesting that these regions are disordered.

To quantify the similarities of the Cɑ positions in the active site and surrounding regions, we aligned the 59 residues that lie within 10 Å of the salicylic acid bound in the active site of SABP2 (Table 7). The similarity of the main chain positions between SABP2 and *Hb*HNL in the 10 Å surrounding the substrate-binding site, 0.62 Å, remains similar to that when comparing the entire proteins, 0.64 Å, Table 6 above. The alignment of the Cɑ positions in the 10-Å region surrounding the substrate-binding site is closer between SABP2 and HNL40 (0.38 Å) and HNL71 (0.28 Å) than it is overall: 0.51 Å and 0.41 Å, respectively. Conversely, the Cɑ positions between *Hb*HNL and the variants in the 10-Å region are more different: HNL40 (0.60 Å) and HNL71 (0.56 Å) versus 0.40 and 0.47 Å overall, respectively. A comparison of the changes between *Hb*HNL and HNL40 and HNL71 show that the Cɑ atoms in this region by have moved by an average of ∼0.6 Å. An alignment of three x-ray crystal structures of SABP2, show an RMSD of 0.24±0.07 Å in this region, so the positions of the Cɑ atoms in HNL71, but not HNL40, are indistinguishable from different structures of SABP2.

**Table 7.**
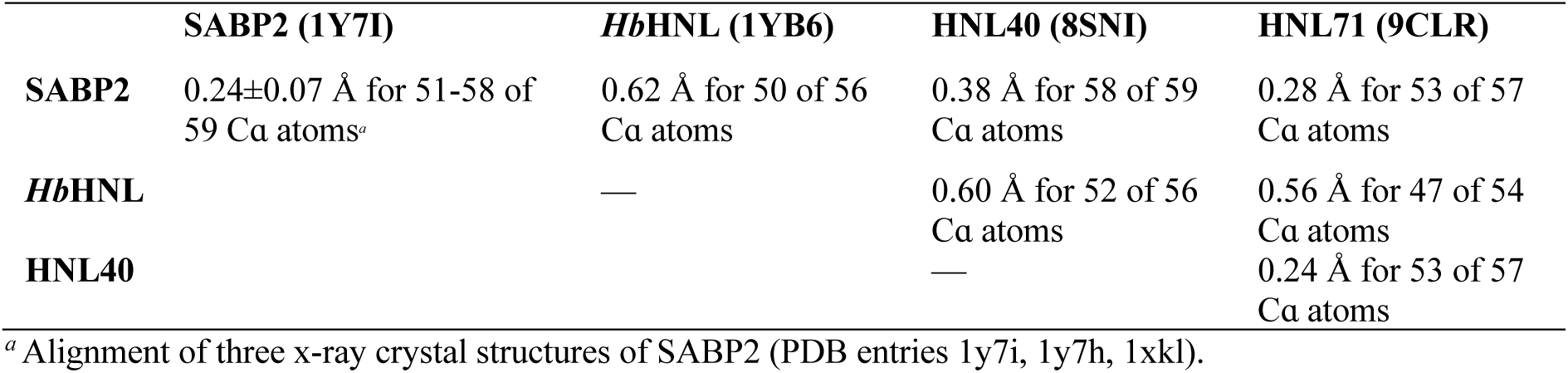
RMSD values for the alignment of Cɑ atoms of the 59 residues within 10 Å of salicylic acid in SABP2.

### 3.5. More distant regions

Another structural change introduced by the changes in HNL40 and HNL71 is the creation of additional tunnels from the active site to the protein surface. This change makes the pattern of tunnels in HNL40 and HNL71 intermediate between those in *Hb*HNL and SABP2 (Fig. 6). *Hb*HNL contains two tunnels, both of which lead to the left in the orientation shown in Fig. 6. In contrast, SABP2 contains seven tunnels. Six of these tunnels lead to the left and one leads to the right. HNL40 and HNL71 each contain four tunnels, three of which lead to the left and one that leads to the right.

**Figure 6.**
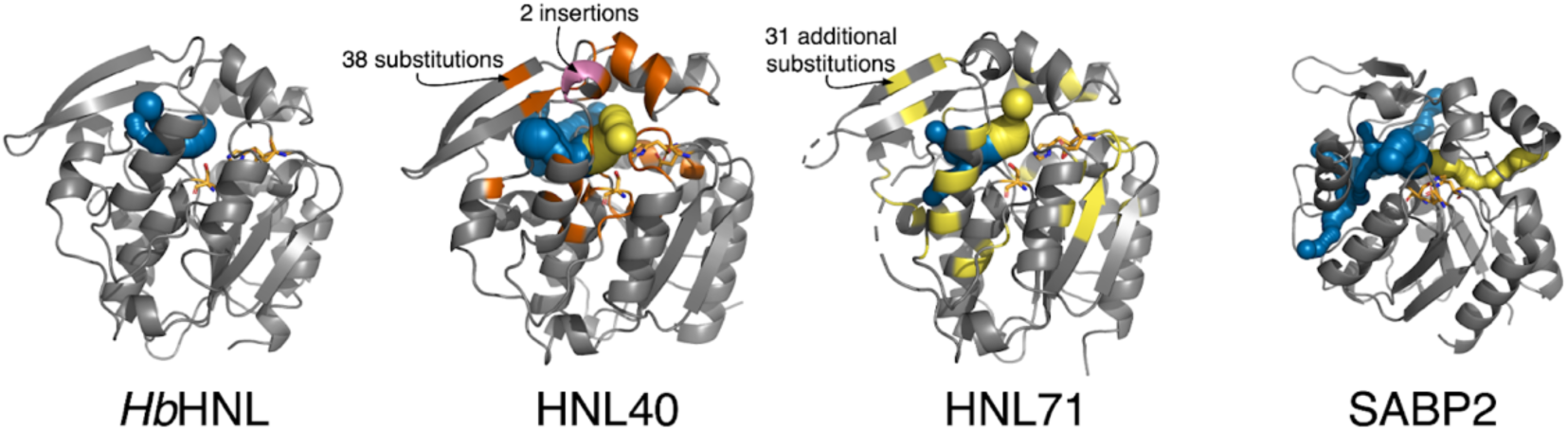
Substitutions in HNL40 and HNL71 create an SABP2-like tunnel (yellow) that is missing in *Hb*HNL. *Hb*HNL (far left) contains two tunnels (blue) that lead from the active site to the left. The catalytic triad is shown as sticks with orange carbons. In contrast, SABP2 (far right) contains seven tunnels, six of these tunnels (blue) lead from the active site to the left, while one tunnel (yellow) leads from the active site to the right. HNL40 and HNL71 each contain four tunnels; three of these tunnels (blue) lead from the active site to the left, while one tunnel (yellow, near the insertion site) leads from the active site to the right. The 39 substitutions in HNL40 are colored orange, while the two insertions are colored pink. The additional thirty-one changes in HNL71 are shown in yellow. Tunnels were identified using Caver Web (Stourac *et al*., 2019) with default settings.

The new tunnel to the right appears near the insertion site (after residue Pro126) of two amino acids in HNL40. The secondary structure of the main chain in this region changes from a loop to a helix for the two inserted residues (Ala127-Glu128) and the two subsequent residues Asn129-Trp130. SABP2 also contains a helix in this region. While this new tunnel to the right in HNL40 and HNL71 creates a tunnel where none existed in *Hb*HNL, the tunnel differs from the corresponding tunnel in SABP2. In HNL40 and HNL71 the tunnel is short and exits to the front in the view shown in Fig. 6. In contrast, in SABP2 the tunnel is longer and exits to the back and right.

### 3.6. Sulfenic acid at position 163 in HNL71

SABP2, *Hb*HNL, HNL71 and HNL40 all contain a cysteine at the position corresponding to 164 in SABP2. During refinement of HNL71, the *F_O_ – F_C_* (red/green) electron density map revealed a notable density adjacent to the sulfur of Cys163 in both chains A and B. Placing an oxygen atom into that density decreased the R_work_ and R_free_ values. This addition of an oxygen atom suggests that Cys163 has been oxidized to a sulfenic acid in HNL71 (Fig. 7). The structures of SABP2, *Hb*HNL and HNL40 show no evidence for oxidation of the corresponding cysteine.

**Figure 7.**
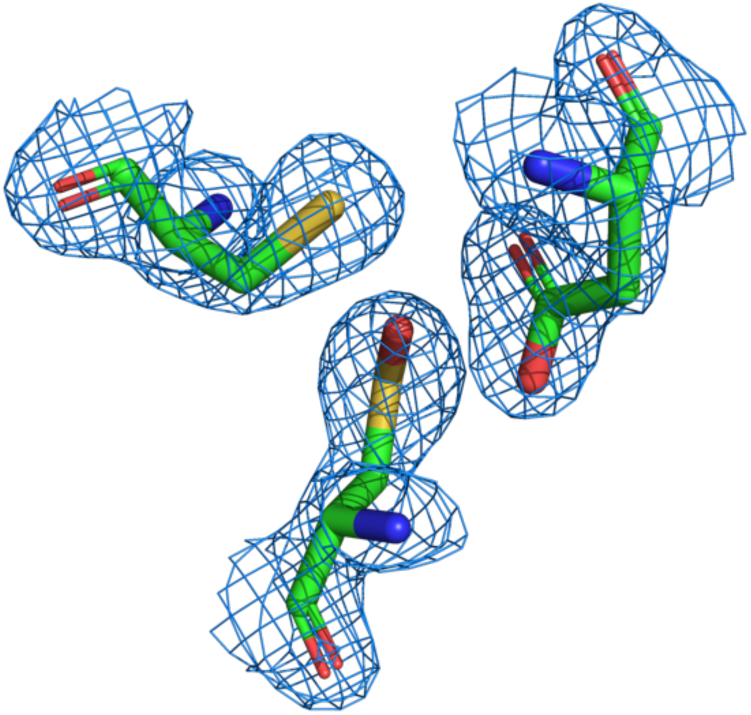
Electron density map showing the sulfenic acid Cso163 (bottom), Asp236 (top right), and Cys242 (top left) in HNL71. The 2*F_O_ – F_C_* map (blue mesh) is contoured at 1.5σ. No oxidation of the corresponding cysteine was observed in the structures of SABP2, *Hb*HNL or HNL40.

The propensity of a cysteine residue to oxidation is sometimes correlated with the nature of nearby amino acids (Garrido Ruiz *et al*., 2022; Salsbury *et al*., 2008), but in the case of HNL71, no oxidation-promoting nearby residue could be identified. The closest residue, Asp236, also occurs in SABP2, *Hb*HNL and HNL40. Comparison of the amino acids within 5 Å of the sulfenic acid at position 163 in HNL71 with the corresponding amino acids in SABP2, *Hb*HNL, and HNL40, where a similar oxidation was not observed, did not identify an amino acid present in HNL71 that was absent from the other three (Table S3). The oxidation seen in HNL71 may be due to oxidative conditions encountered by the HNL71 crystal.

## 4. Conclusions

Despite the 44% sequence identity of hydroxynitrile lyases *Hb*HNL with esterase SABP2, common Ser-His-Asp catalytic triads and α/β-hydrolase protein folds, the two enzymes differ in subtly in structure in the protein core and significantly in the pattern of tunnels and in several regions on the protein surface. The Cα positions in the overall enzymes differ by 0.64 Å. The Cα positions of the catalytic triad and oxyanion hole residues differ by slightly more than that amount, an average of 0.8±0.4 Å, with the largest difference, 1.3 Å, for the oxyanion hole residue OX1. In addition, several residues near the catalytic histidine and aspartate differ by larger amounts, ∼2 Å. In four regions on the enzyme surface the Cα positions differ by large amounts. In the cap domain, the 140-145 loop differs by 11 Å and another loop near residue 129, where SABP2 contains a two-amino-acid insertion, differs by 5.6 Å. In the catalytic domains, the 189-191 surface loop differs by ∼3 Å and the N- and C-terminii differ by 6 and 3 Å, respectively.

Replacing amino acid residues surrounding the substrate-binding region in *Hb*HNL with the corresponding residues from SABP2 created proteins increasingly similar to SABP2. HNL40 shows 59% sequence identity and HNL71 shows 71% sequence identity. In HNL40 all amino acid residues within at least 10 Å of the substrate-binding site are identical to those in SABP2 and in HNL71 all amino acid residues within 14 Å are identical.

The x-ray structures of HNL40 and HNL71 reveal that their core regions are more similar to those of SABP2 and less similar to those of *Hb*HNL. The main chain Cα positions in the 10-Å region surrounding the active site shifted by an average of 0.6 Å to locations where the root-mean-square deviations from the positions in SABP2 were 0.38 Å for HNL40 and 0.28 Å for HNL71. A comparison of three different x-ray structures of SABP2 showed a root-mean-square deviation in the 10-Å region surrounding the active site of 0.24 Å. Since both different SABP2 structures and SABP2 versus HNL71 show similar average Cα atom deviations, the structure of HNL71 in the 10-Å region surrounding the substrate-binding site matches that of SABP2. The average catalytic atom deviation in HNL40 is 0.8 Å, which is larger than the 0.3 Å deviation between three different SABP2 structures, but in HNL71 the average deviation is 0.3 Å making these positions comparable to those in different SABP2 structures. The bound proline in HNL40 binds in a similar position as salicylic acid does in the structure of SABP2. The difference in hydrogen bonding patterns between the two structures is likely due to the ammonium group in the proline and which is lacking in salicylic acid.

The sequence of one of the four regions on the surface of HbHNL that differs from SABP2 is identical in amino acid sequence in HNL40 and HNL71 because it lies close to the substrate binding site. The amino acid residues in the surface loop near residue 129 are identical to those in SABP2 in HNL40 and the conformations are also identical. In HNL71, some of the adjacent loops are also identical in amino acid sequence. The other three surface regions of HNL40 and HNL71 lie more than 14 Å from the substrate-binding region and their amino acid sequences still differ from that in SABP2: the 140-145 loop in the cap domain, the 189-191 surface loop in the catalytic domain and the N- and C-termini. For HNL71, these two surface loops are unresolved suggesting a mixture of conformation in those regions.

Tunnels connect the substrate binding region to the enzyme surface so they involve regions where the amino acid residues differ and where they are identical. *Hb*HNL contains only two tunnels, both exiting between the cap and catalytic domains in the same region, while SABP2 contains seven tunnels most of which exit between the cap and catalytic domains, but also several that pass through the catalytic domain. Both HNL40 and HNL71 contain four tunnels, which all exit between the cap and catalytic domains. A notable new tunnel exits near the 129 loop where the two-amino-acid insertion makes the surface region identical to that in SABP2. The other tunnels in SABP2 involve surface regions that differ from those in HNL40 and HNL71, so they remain absent in these proteins.

Overall, the amino acid substitutions in HNL40 and HNL71 make the core of the enzymes within 10 Å of the substrate binding site and including the positions of the catalytic atoms similar to those in SABP2 for HNL40 and nearly identical for HNL71. One of the surface regions - the lid region near residue 129 - is also identical in amino acid sequence to SABP2, so its structure is also similar to SABP2 in HNL40 and HNL71. Three other surface regions differ in SABP2 and *Hb*HNL and retain both their amino acid residue differences and structure differences in HNL40 and HNL71. Since tunnels involve surface residues, these remaining differences on the surface account for differences in the tunnels between SABP2 and HNL40 and HNL71.

## Supporting information

**Figure S1.**
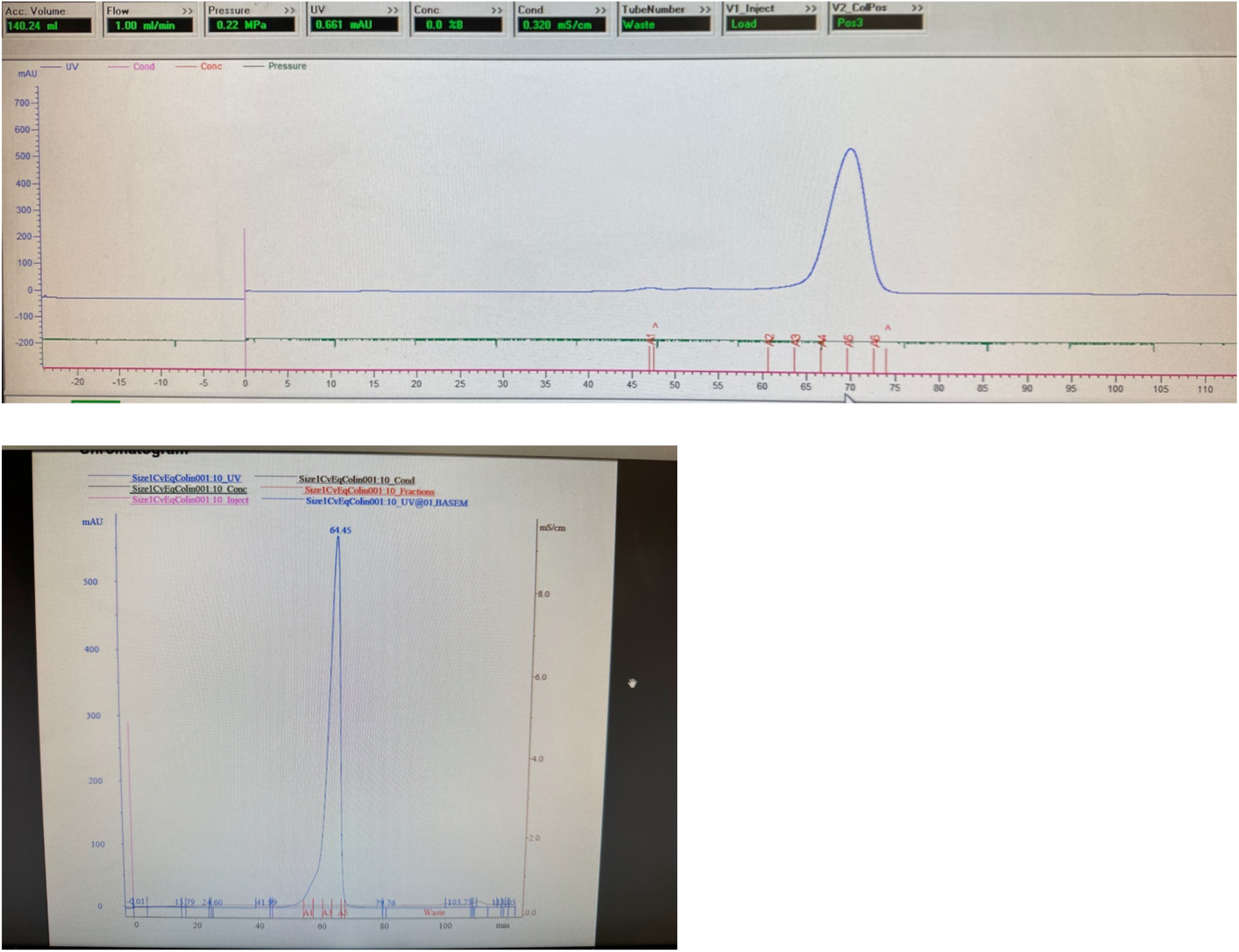
Size-exclusion chromatography traces of HNL71 purification from February 8, 2023 (top) and HNL40 purification from March 19, 2023 (bottom) indicate high purity. UV absorbance at 280 nm (blue trace), given in milli-absorbance units (mAU), indicates the presence of protein eluting as single, symmetrical peaks at ∼70 minutes (HNL71) and ∼64 minutes (HNL40), consistent with previously observed elution times for *Hb*HNL-based variants. The absence of additional peaks suggests minimal aggregation or degradation of the protein samples.

**Figure S2.**
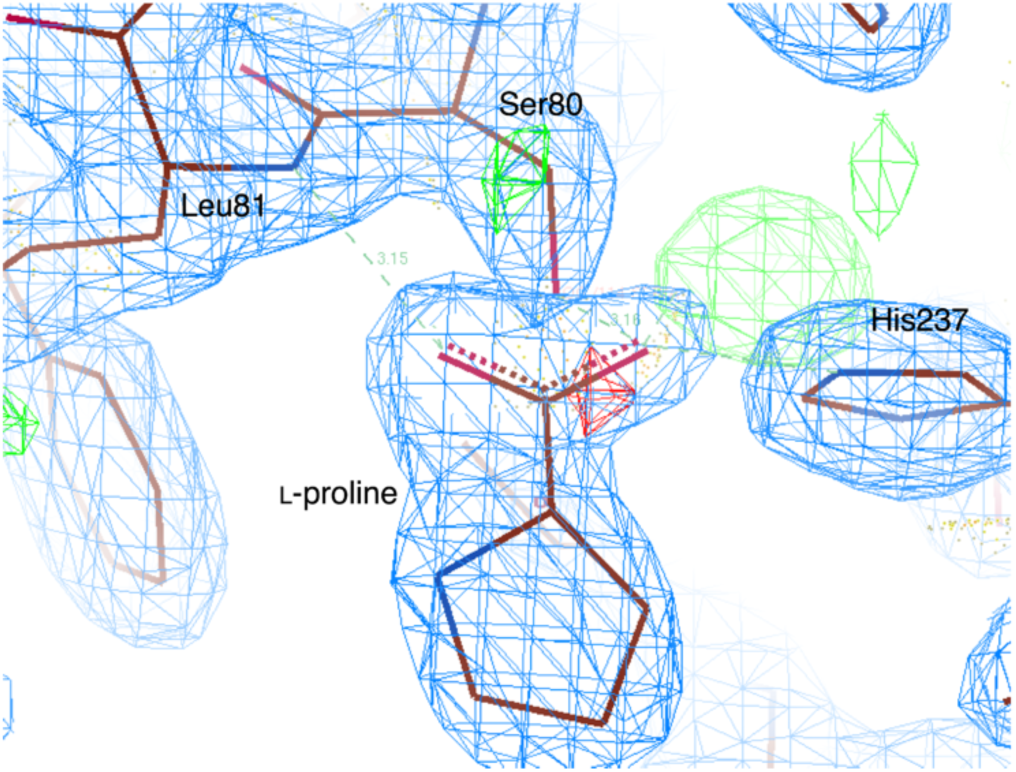
The active site of the HNL40 structure contains a bound L-proline which originates from the crystallization solution and mimics a bound product carboxylate of ester hydrolysis. The electron density map (2*F_O_ – F_C_* (blue mesh) contoured at 1.5σ) and stick representation of the bound L-proline shows some of the hydrogen bonds that the carboxylate oxygens of the proline accept from the catalytic residues. One carboxylate oxygen of the bound proline accepts hydrogen bonds from the oxyanion hole N-H (Ala12 d_N-O_ = 2.76 Å (not shown), Leu81 d_N-O_ = 3.15 Å), while the other carboxylate oxygen accepts hydrogen bonds from the Oɣ-H of the catalytic serine 80 (d_O-O_ = 3.16 Å), Nε-H of the catalytic histidine 237 (d_O-N_ = 2.87 Å) and a water molecule 416 (d_O-O_ = 2.68 Å; not shown).

**Table S1.**
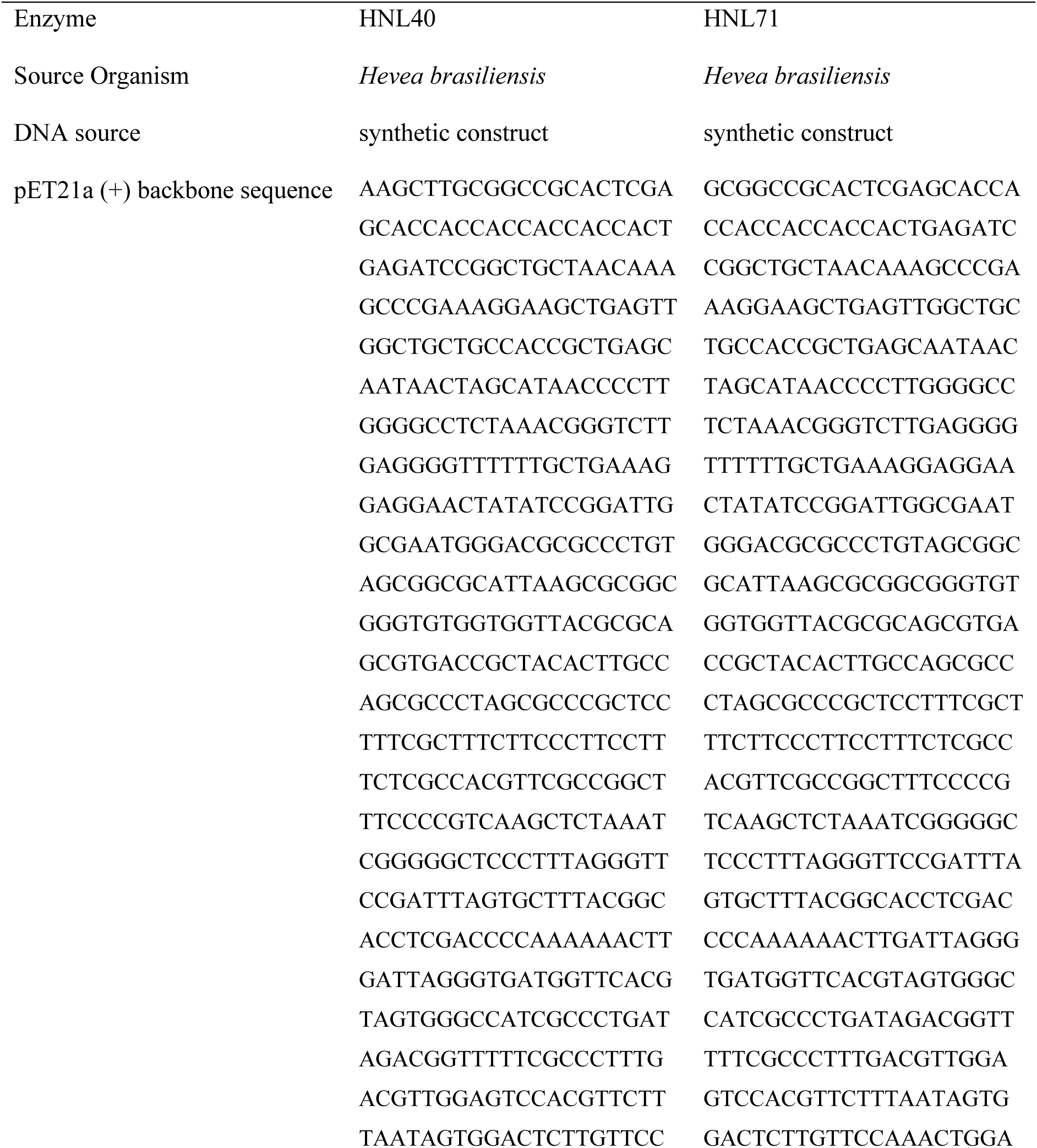
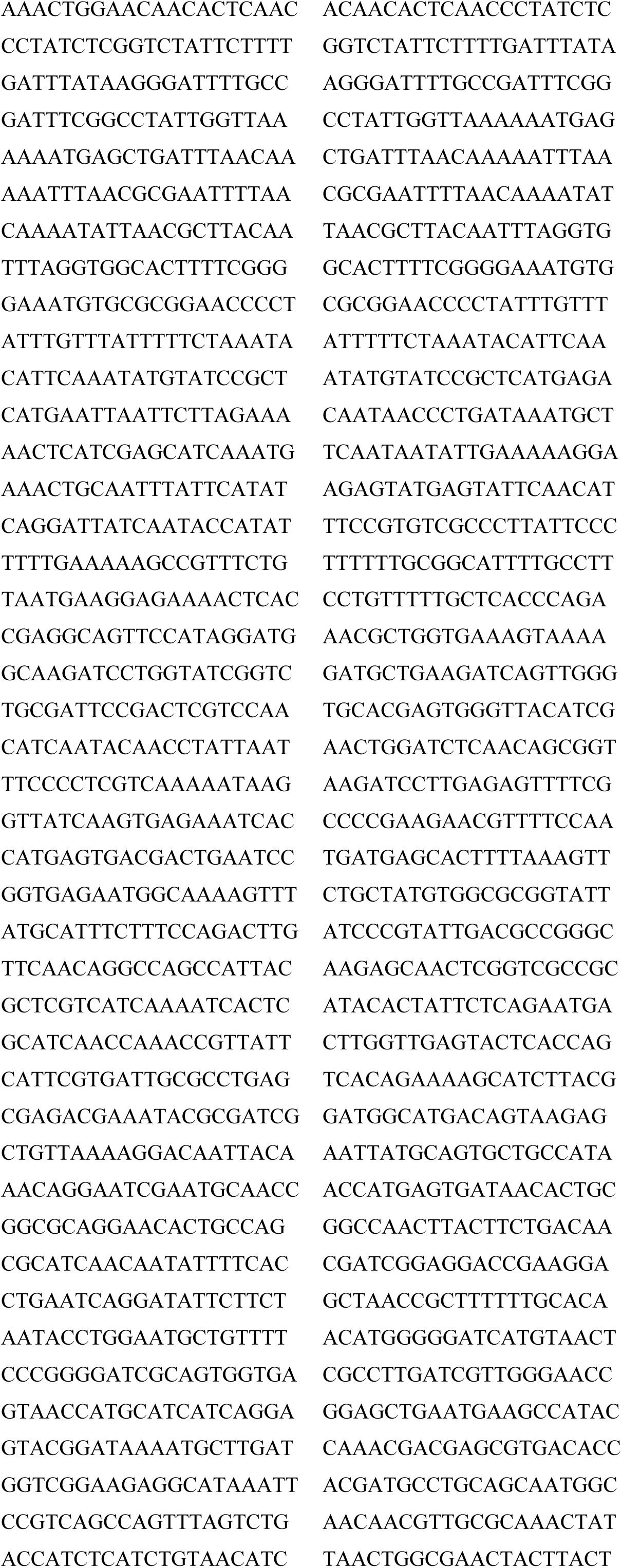
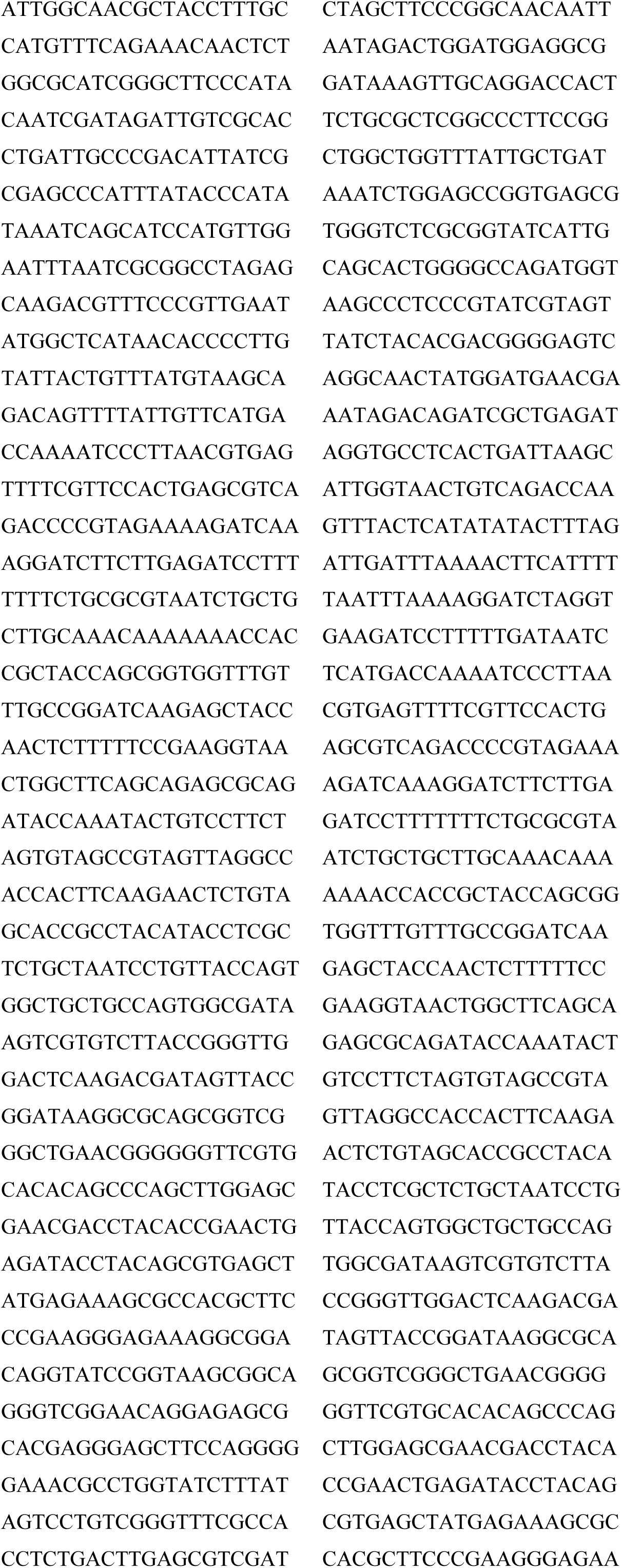
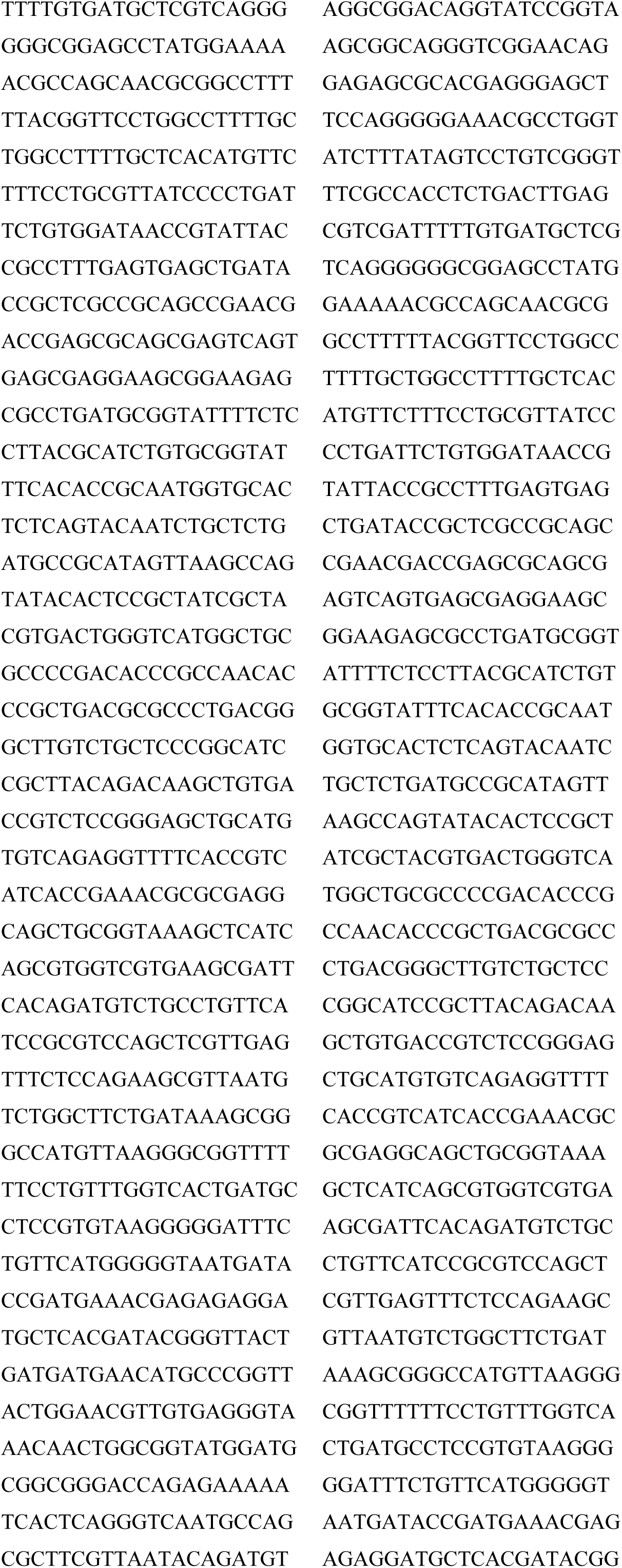
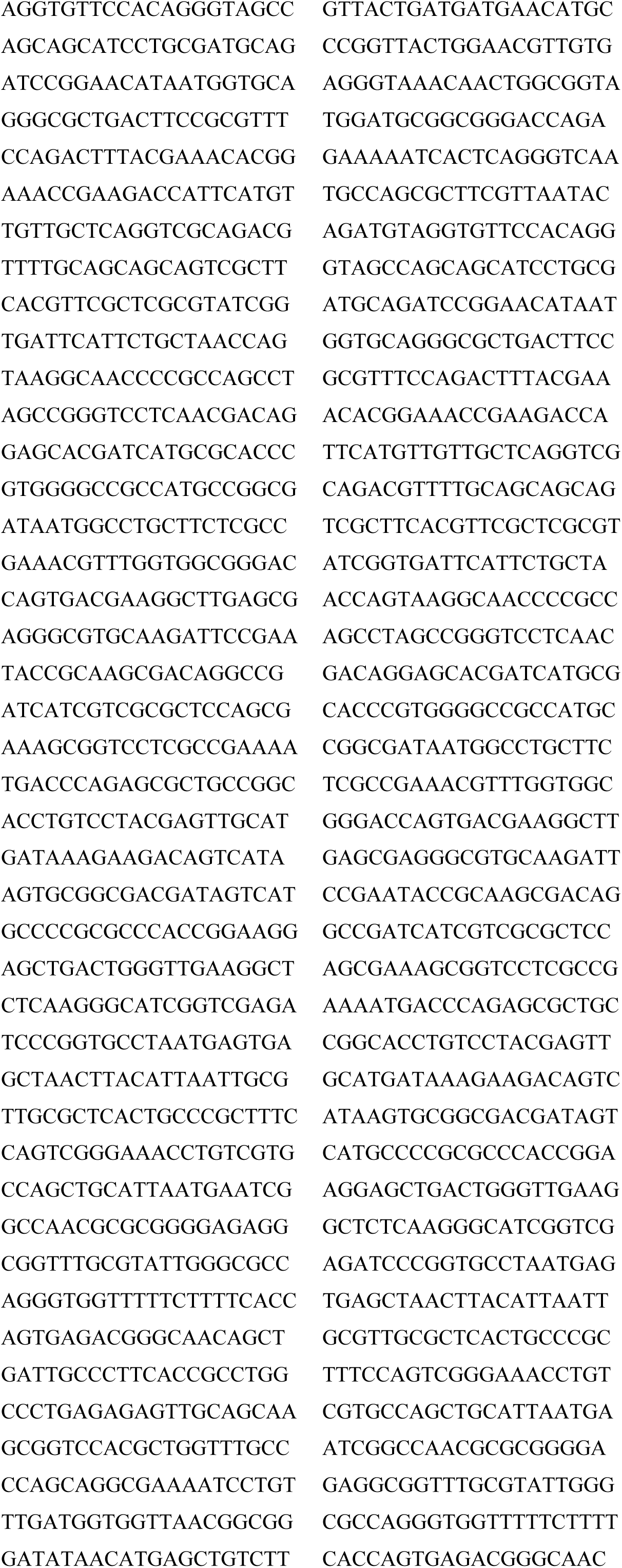
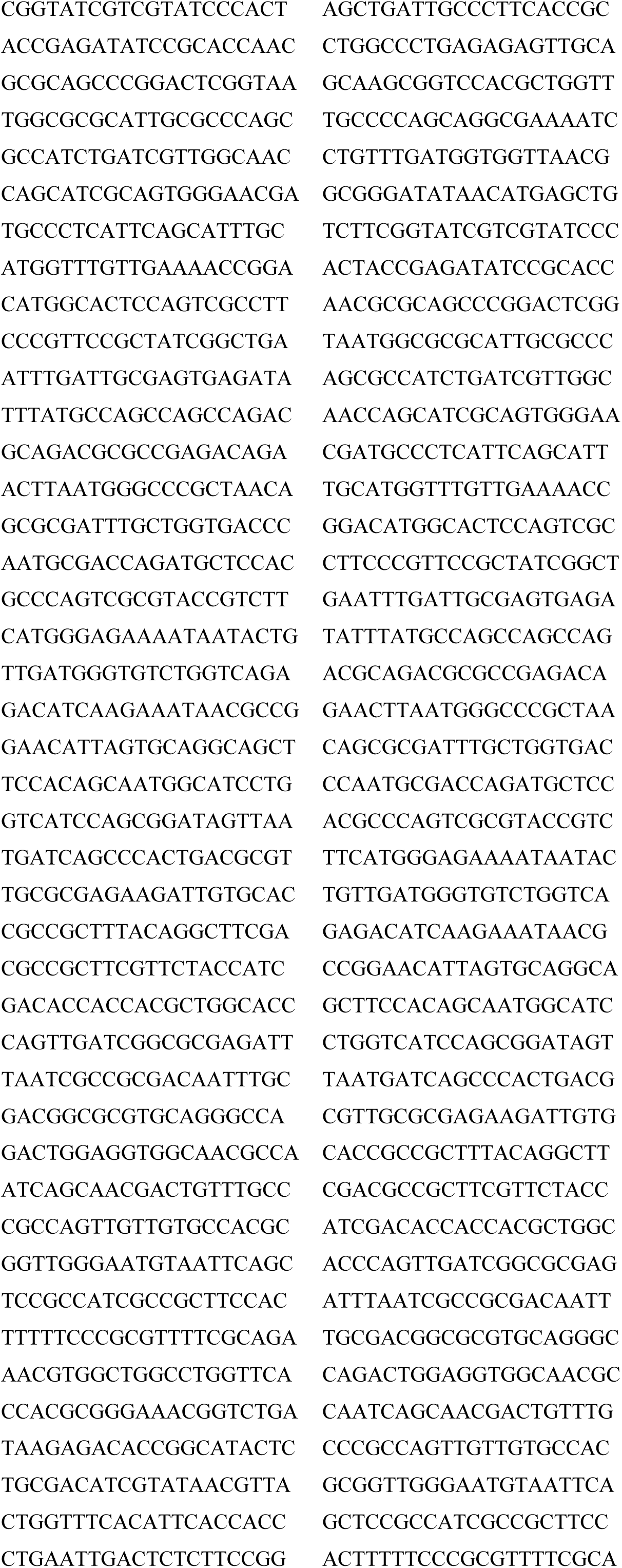
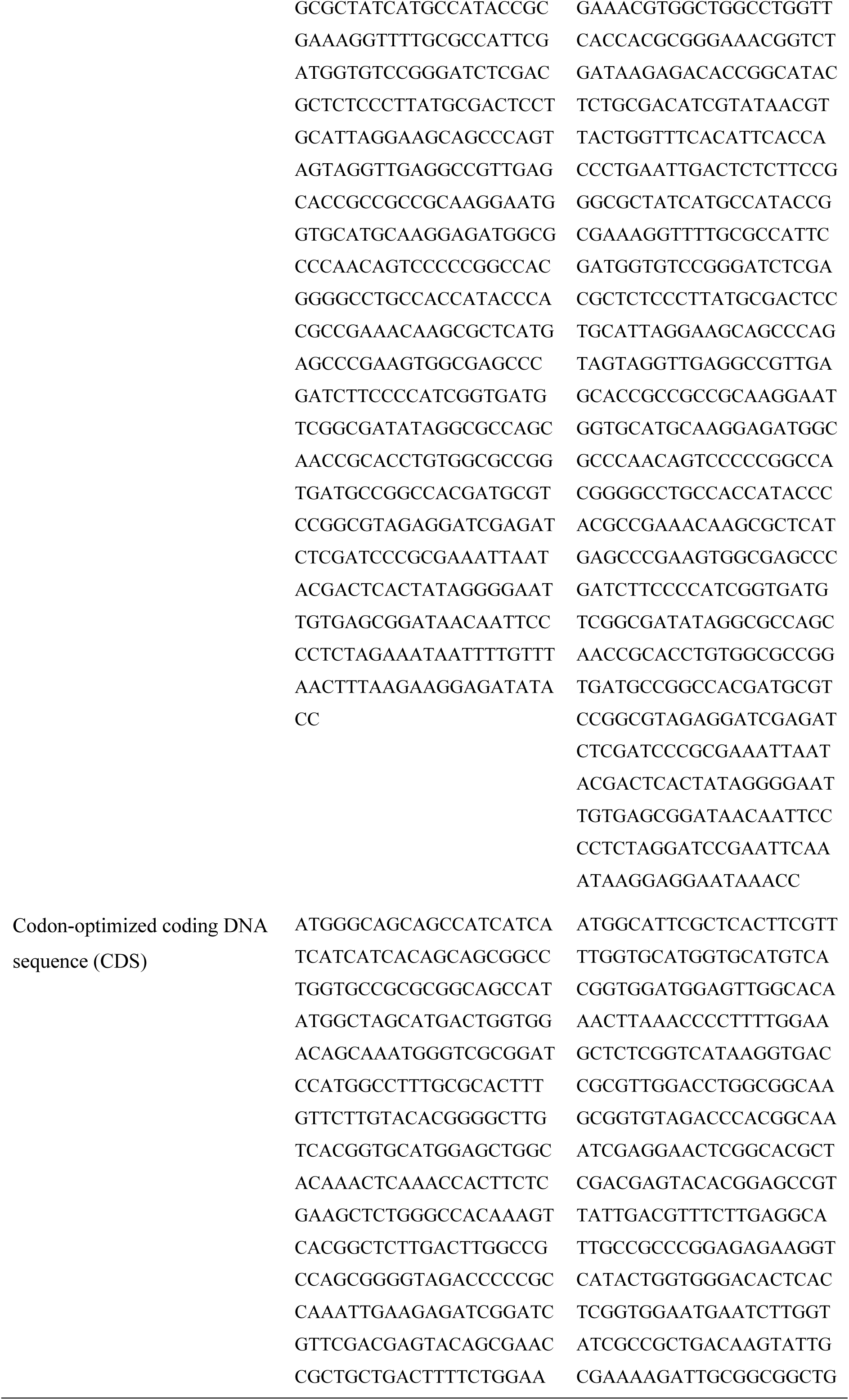
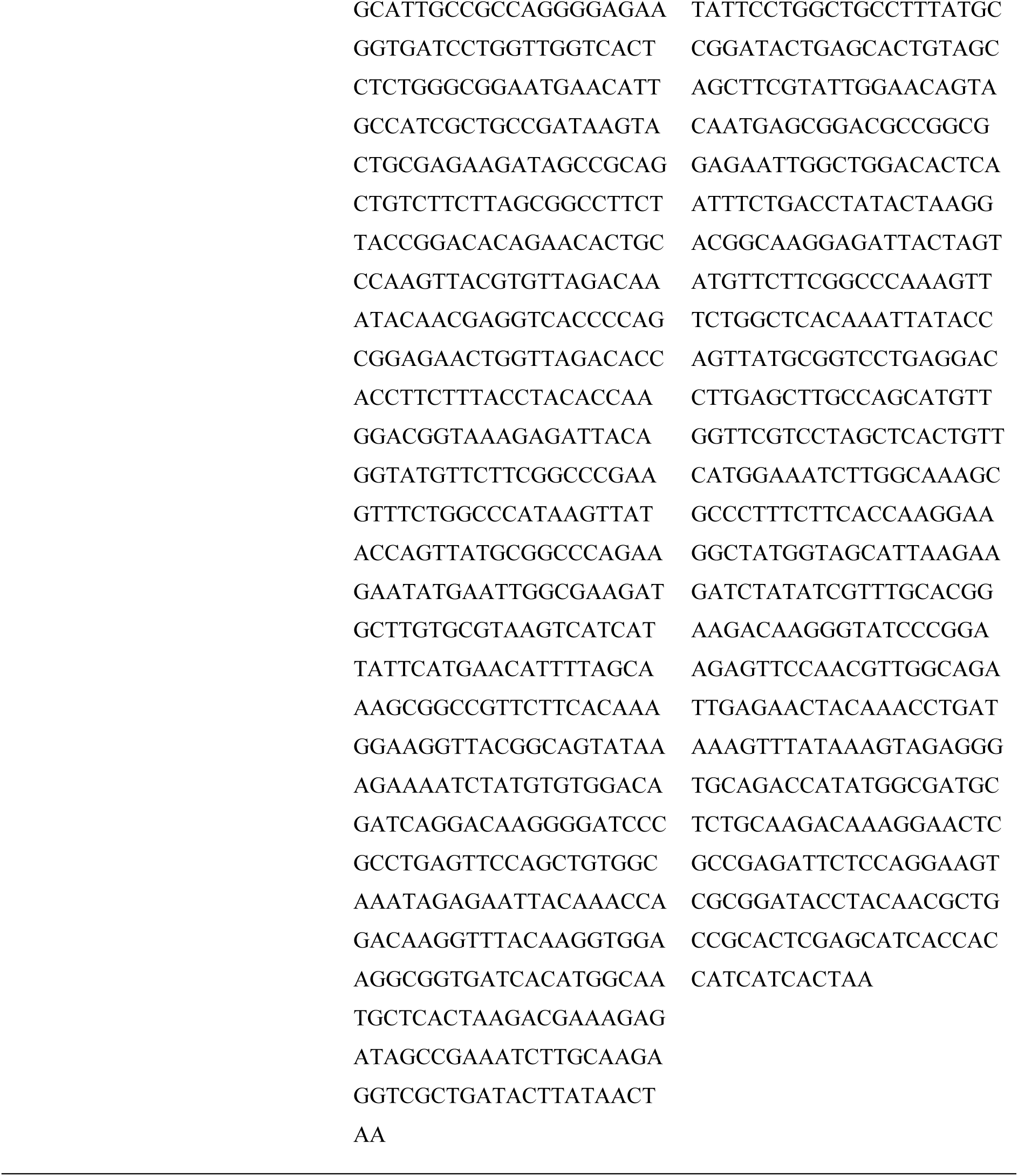
DNA sequence of the protein expression plasmids used to make HNL40 and HNL71.

**Table S2.**
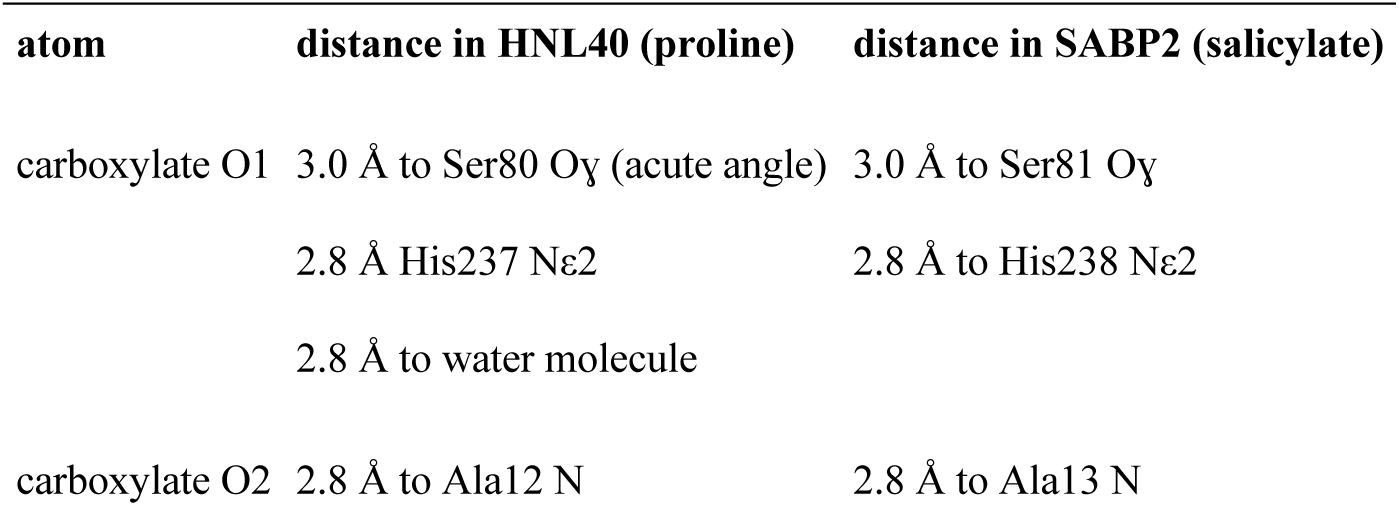
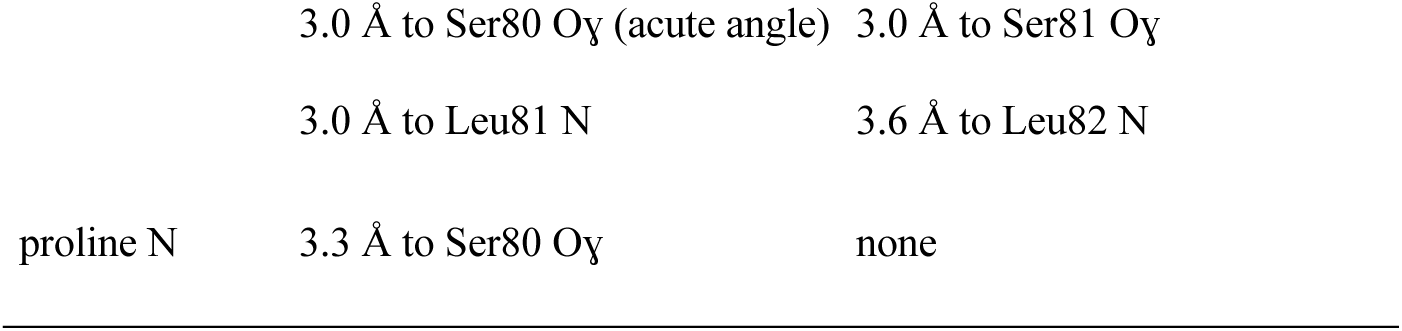
Distances between the polar atoms in HNL40 (8sni) and SABP2 (1y7i) and a bound carboxylate-containing molecule, proline or salicylate, respectively.

| atom | distance in HNL40 (proline) | distance in SABP2 (salicylate) |
| --- | --- | --- |
| carboxylate O1 | 3.0 Å to Ser80 Oy (acute angle) | 3.0 Å to Ser81 Oy |
|  | 2.8 Å His237 Nε2 | 2.8 Å to His238 Nε2 |
|  | 2.8 Å to water molecule |  |
| carboxylate O2 | 2.8 Å to Ala12 N | 2.8 Å to Ala13 N |
| | 3.0 Å to Ser80 Oγ (acute angle) | 3.0 Å to Ser81 Oγ |
|  | 3.0 Å to Leu81 N | 3.6 Å to Leu82 N |
| proline N | 3.3 Å to Ser80 Oγ | none |

**Table S3.** Residues near CSO163 in HNL71 and the corresponding residues in SABP2, HNL40 and *Hb*HNL cannot account for the oxidation in HNL71, but not the other three proteins. Eight amino acid residues lie with at least one atom within 5 Å of Sɣ of residue 163 in HNL71. Four of these residues (Tyr160, Leu162, Asp236, Leu241) are identical to the corresponding residues in SABP2, *Hb*HNL and HNL40. A comparison of the remaining four residues shows that at least one of the non-oxidized proteins contains the same amino acid as the one present in HNL71. This lack of a unique nearby amino acid suggests that the oxidation was due to oxidative conditions encountered by the HNL71 crystal and not due to the enhanced reactivity of Cys163 in HNL71.

| HNL71 | SABP2 | HNL40 | *HbHNL* |
| --- | --- | --- | --- |
| Gly164 | Ser165 | Gly164 | Gly162 |
| Asp167 | Asp168 | Glu167 | Glu165 |
| Met238 | Met239 | Met238 | Lys236 |
| Cys242 | Cys243 | Thr242 | Thr240 |

